# The Parkinson’s disease risk gene cathepsin B promotes fibrillar alpha-synuclein clearance, lysosomal function and glucocerebrosidase activity in dopaminergic neurons

**DOI:** 10.1101/2023.11.11.566693

**Authors:** Jace Jones-Tabah, Kathy He, Konstantin Senkevich, Nathan Karpilovsky, Ghislaine Deyab, Yuting Cousineau, Daria Nikanorova, Taylor Goldsmith, Esther del Cid Pellitero, Carol X-Q. Chen, Wen Luo, Zhipeng You, Narges Abdian, Isabella Pietrantonio, Thomas Goiran, Jamil Ahmad, Jennifer A. Ruskey, Farnaz Asayesh, Dan Spiegelman, Cheryl Waters, Oury Monchi, Yves Dauvilliers, Nicolas Dupré, Irina Miliukhina, Alla Timofeeva, Anton Emelyanov, Sofya Pchelina, Lior Greenbaum, Sharon Hassin-Baer, Roy N. Alcalay, Austen Milnerwood, Thomas M. Durcan, Ziv Gan-Or, Edward A. Fon

## Abstract

Variants in the *CTSB* gene encoding the lysosomal hydrolase cathepsin B (catB) are associated with increased risk of Parkinson’s disease (PD). However, neither the specific *CTSB* variants driving these associations nor the functional pathways that link catB to PD pathogenesis have been characterized. CatB activity contributes to lysosomal protein degradation and regulates signaling processes involved in autophagy and lysosome biogenesis. Previous *in vitro* studies have found that catB can cleave monomeric and fibrillar alpha-synuclein, a key protein involved in the pathogenesis of PD that accumulates in the brains of PD patients. However, truncated synuclein isoforms generated by catB cleavage have an increased propensity to aggregate. Thus, catB activity could potentially contribute to lysosomal degradation and clearance of pathogenic alpha synuclein from the cell, but also has the potential of enhancing synuclein pathology by generating aggregation-prone truncations. Therefore, the mechanisms linking catB to PD pathophysiology remain to be clarified. Here, we conducted genetic analyses of the association between common and rare *CTSB* variants and risk of PD. We then used genetic and pharmacological approaches to manipulate catB expression and function in cell lines and induced pluripotent stem cell-derived dopaminergic neurons and assessed lysosomal activity and the handling of aggregated synuclein fibrils. We find that catB inhibition impairs autophagy, reduces glucocerebrosidase (encoded by *GBA1*) activity, and leads to an accumulation of lysosomal content. In cell lines, reduction of *CTSB* gene expression impairs the degradation of pre-formed alpha-synuclein fibrils, whereas *CTSB* gene activation enhances fibril clearance. In midbrain organoids and dopaminergic neurons treated with alpha-synuclein fibrils, catB inhibition potentiates the formation of inclusions which stain positively for phosphorylated alpha-synuclein. These results indicate that the reduction of catB function negatively impacts lysosomal pathways associated with PD pathogenesis, while conversely catB activation could promote the clearance of pathogenic alpha-synuclein.

## Introduction

Parkinson’s disease (PD) is characterized by both the degeneration of dopaminergic neurons in the substantia nigra and by the accumulation of Lewy bodies; proteinaceous inclusions composed in largely of misfolded and aggregated α-synuclein (α-syn) ^1^. Mutations that increase protein levels of α-syn or its propensity to aggregate contribute substantial genetic risk to PD ^2,3^, supporting the predominant hypothesis that α-syn aggregation is a key step in the pathological cascade leading to neurodegeneration in PD. The lysosome serves as the principal site for degradation of aggregated α-syn ^4–6^, and mutations in lysosomal genes also represent a substantial genetic risk for PD ^7^. Thus, there is great interest in understanding the lysosomal pathways that mediate α-syn clearance. Cathepsin B (catB, encoded by the *CTSB* gene) is a proteolytic enzyme of the cysteine cathepsin family with endo– and exo-peptidase activity that is normally localized to the lysosomal lumen ^8^. CatB has been implicated both in the lysosomal degradation of α-syn and as a genetic risk factor for PD. In the present study we further elucidate the relationship between *CTSB* variants and PD risk and demonstrate that catB modulates lysosome function and the clearance of α-syn aggregates in cell lines and human dopaminergic neurons.

The importance of lysosomal function in PD is well established by both functional and genetic studies ^7^. Recently, genome-wide association studies (GWAS) have identified significant association between variants in the *CTSB* genetic locus and the risk of PD generally ^9^ and specifically in carriers of pathogenic *GBA1* variants ^10^. In addition to genetic evidence linking *CTSB* to PD, catB protein or activity levels are reduced in several cellular models of PD. For example, pathological α-syn species have been shown to impair catB trafficking to the lysosome ^11^, while iPSC derived neurons harboring mutations in *SNCA* or *GBA1* exhibited reduced catB activity ^12,13^. Additionally, knockout of the PD risk gene *TMEM175* impairs catB activity by destabilizing lysosome pH ^14,15^, while mutations in *LRRK2,* the most common cause of familial PD, have been shown to suppress catB expression or activity in the lysosome ^16,17^. Thus, several lines of evidence suggest that disrupted catB function could play a role in PD pathogenesis.

One potential mechanism linking catB to PD is through its ability to cleave both monomeric and aggregated forms of α-syn, which has been demonstrated *in vitro* ^18–20^. However, while this could argue for a protective role of catB against synucleinopathy, the α-syn truncations produced by *in vitro* catB cleavage exhibit an increased propensity to aggregate ^21^ and although lysosome function is essential for degradation of fibrillar α-syn ^22^, it has also been suggested that catB activity contributes to α-syn toxicity in some cellular models ^23^. Moreover, catB has been linked to the α-syn dependent activation of inflammatory pathways ^24^ and is a key regulator of cell death in many cellular contexts ^25^. Thus, there are compelling arguments to be made in favor of both protective and potentially pathogenic actions of catB in the etiology of PD and its specific role remains to be elucidated.

Here, we aim to both clarify the genetic evidence pertaining to how *CTSB* variants may contribute to PD etiology, and to functionally characterize the role of catB in relation to lysosome function and α-syn clearance. We first provide genetic evidence that PD-associated *CTSB* variants decrease expression levels of the enzyme. Second, by pharmacologically and genetically modulating catB expression or activity in cell lines and human dopaminergic neurons we demonstrate that catB is required for lysosomal functions including glucocerebrosidase activity and contributes to clearance of fibrillar α-syn. These findings argue in favor of a protective effect of catB in PD.

## Materials and Methods

### Fine mapping and eQTL analysis of *CTSB* variants

To identify the most likely variant driving the PD-association in the *CTSB* locus, we performed analyses using the summary statistics from the most recent PD GWAS ^9^ and multiple bioinformatic tools. First, to examine whether there are multiple independent associations in this locus, we used genome-wide complex trait conditional and joint analysis (GCTA-COJO) ^26^, using default parameters. For downstream analyses, we generated a linkage disequilibrium (LD) matrix for the *CTSB* locus using PLINK 1.9 ^27^, including all variants within ± 1Mbp from the top variant. Then, we performed fine-mapping of the *CTSB* locus to nominate the most likely driving variants using FINEMAP ^28^, with minor allele frequency (MAF) threshold > 0.01. Expression quantitative trait locus (eQTL) analysis was performed using colocalization (COLOC) ^29^, which examines whether the same variants associated with the trait (PD) are also associated with gene expression. QTLs were tested in a total of 109 tissues and cells (Supplementary Table 1) To further explore the link between genetic variants, QTLs and PD we used Summary-data-based Mendelian Randomization (SMR), which uses Mendelian randomization to suggest potential causality, followed by heterogeneity in dependent instruments (HEIDI) to distinguish between pleiotropy (or causality) and LD ^30^.

### Rare Variant Analysis

Rare variant analysis was performed on 5,801 PD cases and 20,427 controls across six cohorts (Supplementary Table 2). All patients were diagnosed by movement disorder specialists according to the UK brain bank criteria ^31^ or MDS diagnostic criteria ^32^. From the AMP-PD and UKBB cohorts we only included participants of European ancestry and excluded any first and second-degree relatives from the analysis. Quality control procedures for AMP-PD and UKBB were performed as previously described in detail ^33,34^.

In addition, we conducted sequencing on four distinct cohorts at McGill University (McGill cohort, Columbia cohort, Sheba Medical Center cohort and Pavlov and Human Brain Institutes cohort). We performed sequencing of the *CTSB* gene, including exon-intron boundaries (±50bps) and the 5’ and 3’ untranslated regions (UTRs) using molecular inversion probes (MIPs) as described previously ^35^. The full protocol is available at https://github.com/gan-orlab/MIP_protocol. Library sequencing was performed by the Genome Quebec Innovation Centre on the Illumina NovaSeq 6000 SP PE100 platform. We used Genome Analysis Toolkit (GATK, v3.8) for post-alignment quality checking and variant calling ^36^. We applied standard quality control procedures ^37^. In brief, only variants with minor allele frequency (MAF) of less than 1% and a minimum quality score (GQ) of 30 were included in the analysis. The average coverage for *CTSB* in cohorts sequenced at McGill was >4000X with 95% nucleotides covered at 30× (Supplementary Table 3).

To analyze rare variants, we applied the optimized sequence Kernel association test (SKAT-O, R package) ^38^ with further meta-analysis between the cohorts using metaSKAT package ^39^. We performed separate analyses for the whole gene, non-synonymous and functional (nonsynonymous, stop and frameshift variants) and variants with Combined Annotation Dependent Depletion (CADD) scores of ≥ 20 ^40^. We adjusted for sex, age and ethnicity in all analyses. We also analyzed whether rare *CTSB* variants affected age at onset of PD.

### Generation of CTSB-KO and SNCA-KO iPSC

The previously characterized AIW002-02 iPSC cell line ^41^ was used to generate *CTSB* and *SNCA* knockout lines. CRISPR gRNAs were designed using Synthego and the sequences of reagents used are depicted in Supplementary Table 4. The SNCA-KO line was created by using two gRNAs to introduce a 122bp deletion into exon 1 of the gene. The gRNA sequences were cloned into a Cas9/puromycin expression vector PX459 (Addgene, #48139) and transfected into iPSCs using Lipofectamine ^TM^ stem reagent (ThermoFisher Scientific). Transfected iPSCs were selected in 0.3μg/mL puromycin for 72h and surviving colonies were manually picked and expanded for PCR screening to confirm deletion of the target region. Colonies confirmed to be knockout by PCR screening were further validated by sanger sequencing, and loss of protein was confirmed by Western blot in differentiated neurons.

The *CTSB*-KO cell line was created by HDR using a single gRNA and ssDNA repair template to introduce a stop tag in exon 4 of the gene. Cas9 nuclease, gRNA and a ssDNA repair template for HDR were introduced by Lonza Nucleofection. Edited alleles were detected with ddPCR to select edited clones and deletion was verified by PCR screening followed by sanger sequencing, and finally loss of protein was determined by Western blot (Supplementary Fig 1).

All lines were subject to quality control as previously described ^41^ and included verification of pluripotency by immunofluorescent staining for pluripotency markers (Nanog, Tra1-60, SSEA4 and OCT3/4), verification of normal karyotype and verification of normal profile on genome stability test (Supplementary Fig 1).

### iPSC culture and dopaminergic neuron differentiation

All cell culture reagents used, and media compositions are depicted in Supplemental Table 5. Midbrain neuronal precursor cells (NPCs) and dopaminergic neurons were generated following previously established protocols ^42,43^. Briefly, iPSCs were dissociated with Gentle Cell dissociation reagent and transferred to uncoated flasks in NPC Induction Media to allow for embryoid bodies (EB’s) to form. EBs were cultured for 7 days and then transferred to polyornithine/laminin coated flasks and grown for another 7 days in NPC induction media. To expand NPCs the EBs were then dissociated into small colonies by trituration in Gentle Cell dissociation media and replated as a monolayer on polyornithine/laminin coated flasks. After reaching confluence, NPCs were harvested and frozen in FBS with 10% DMSO and stored in liquid nitrogen.

To differentiate neurons, NPCs were thawed in NPC Maintenance Media with Y-27632 (ROCK inhibitor) and plated on polyornithine/laminin. NPCs were grown for 5-9 days until confluent. For final differentiation into dopaminergic neurons, NPCs were dissociated using Accutase and plated on polyornithine/laminin in Dopaminergic Differentiation Media. After 5 days, media was supplemented with mitomycin C to remove proliferative cells. Dopaminergic neurons were maintained by exchanging 1/3 of the culture volume for fresh dopaminergic differentiation media every 5-7 days. Neurons from every batch were assessed by immunofluorescence for expression of Map2 and TH (Supplementary Fig 2A,B), and only batches achieving at least 50% Map2/TH positivity after 4 weeks of differentiation were used for the experiments included in this manuscript.

For high-content imaging experiments neurons were plated on 96-well plates at a density of 15,000 cells per well. For protein and RNA isolation experiments neurons were plated on 6-well plates at a density of 750,000 cells per well. For live-imaging experiments, neurons were plated on 4-chamber imaging dishes at a density of 100,000 cells per well.

### Organoid culture, treatment, and imaging

The patient derived iPSCs with SNCA triplication mutation (3×SNCA) and corresponding SNCA knockout (SNCA-KO) were previously described ^44^ and provided by Dr. Tilo Kunath. These cells were used to generate midbrain organoids following a protocol previously established in our labs ^45,46^. Three months after organoid induction they were treated with either DMSO (vehicle) or 1 μM CA074me and treatment was maintained for 60 days. All organoids (12 per group) were fixed, cryo-sectioned, and prepared for immunofluorescence using antibodies against Map2, TH, α-syn and pSyn-129 as previously described ^45^. Verification of expression of TH and absence of α-syn in SNCA-KO in the organoids used in this study is depicted in Supplementary Figure 3. Images were acquired using the Leica TCS SP8 confocal microscope and image analysis was performed with an in-house developed script for quantification of immunofluorescent signal in organoids (https://github.com/neuroeddu/OrgQ).

### **α**-synuclein preformed fibril (PFF) generation and characterization

PFFs were generated from recombinant α-synuclein monomers as previously described ^47,48^. All PFFs underwent quality control assessment by electron microscopy (Supplementary Fig. 2C,D).

### RPE1 CRISPRa and CRISPRi Cell Line Generation

Human retinal pigment epithelial-1 cells (RPE1) were grown in Dulbecco’s Modified Eagle Medium (Wisent) supplemented with 10% fetal bovine serum (Wisent). To generate CRISPRa and CRISPRi parental cell lines, lentivirus was used to stably transduced RPE1 cells with either pLX_311-KRAB-dCas9 ^49^ (Addgene #96918, henceforth referred to as CRISPRi) or EF1a-FLAG-dCas9-VPR ^50^ (Addgene #114195, henceforth referred to as CRISPRa) and single clones were selected and characterized to generate monoclonal parental lines stably expressing the CRISPRa and CRISPRi machinery. The gRNA sequences targeting our genes of interest were selected from previously published CRISPRa/i libraries ^51^ (Supplemental Table 6), synthesized by IDT and cloned into pCRISPRi/a-v2 ^51^ (Addgene #84832). Lentivirus was used to stably transduce parental CRISPRa and CRISPRi cell lines which then underwent puromycin selection to generate polyclonal cell lines expressing the gRNA of interest. For each target, several gRNAs were tested and the best performing sequences were selected by assessing target modulation by RT-qPCR analysis of gene expression.

### Drug and PFF treatments

CA074me (Selleckchem) and PADK (Bachem) treatments were performed at the indicated final concentrations with DMSO as vehicle. For PFF experiments in Figures 2 and 3, a single drug treatment was performed simultaneous with PFF administration, after which media was refreshed every 5-7 days. For lysosomal assays in Fig 3, drug was administered 5 days prior to the assay unless otherwise specified.

**Figure 1:**
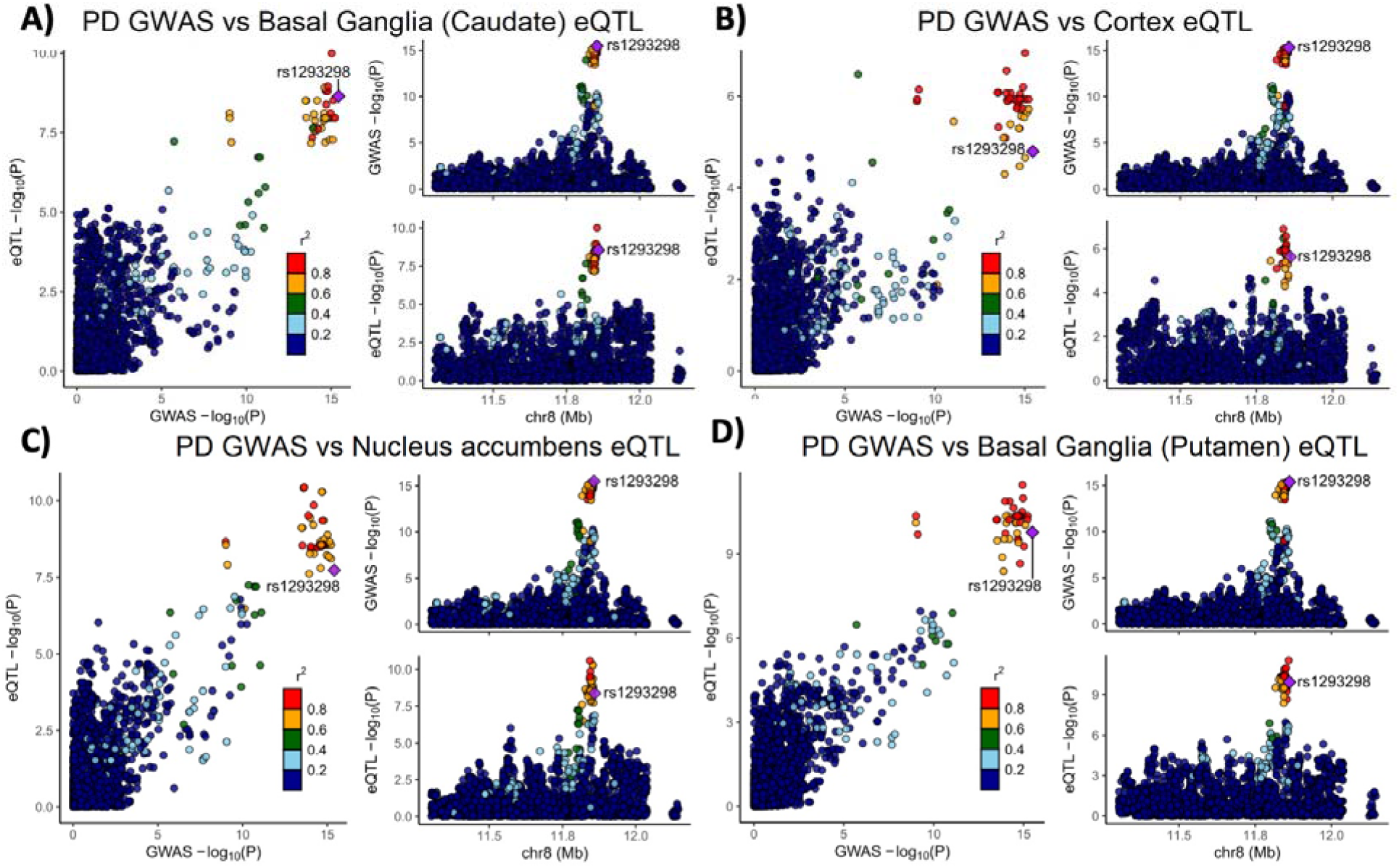
Genetic dissection of the *CTSB* locus in Parkinson’s disease risk. Locus zoom plots depicting the *CTSB* locus (±500 kb) in Parkinson’s disease GWAS with brain eQTLs. The top PD-associated variant (rs1293298) is highlighted in purple, and variants in strong LD (r^2^>0.8) are in red. Each panel includes three plots: The left plot in each panel compares the p values from the PD GWAS and expression data for each variant. Variants that are in the top right corner of this plot are therefore associated with both risk of PD and *CTSB* expression. The top right plots depict the PD GWAS association in this locus and is identical in all four panels. On the bottom right of each panel, the plot depicts the association between variants in this locus and *CTSB* RNA expression in the relevant tissue. (A) PD GWAS plotted together with Basal ganglia (Caudate) eQTL. (B) PD GWAS plotted together with Cortex eQTL. (C) PD GWAS plotted together with Nucleus accumbens eQTL. (D) PD GWAS plotted together with Basal ganglia (Putamen) eQTL.

**Figure 2:**
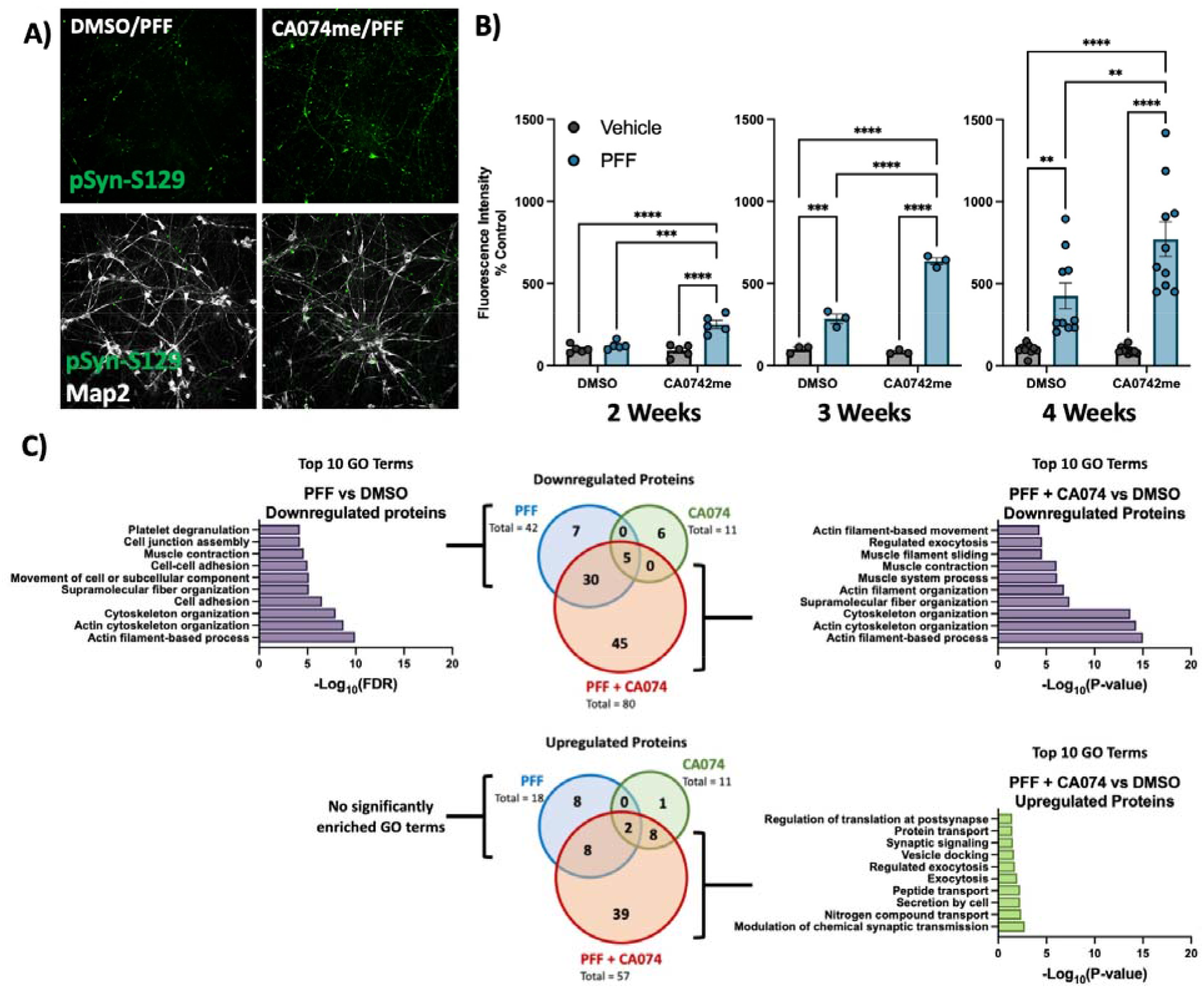
Cathepsin B inhibitors potentiate the effect of α-syn PFFs on dopaminergic neurons: A) Representative immunofluorescent images from high-content confocal imaging of DA neurons treated with CA074me (1 μM) and/or α-syn PFFs (300 nM) and stained for Map2 and pSyn-S129. B) Quantification of pSyn-S129 in Map2-positive cells 2, 3, or 4 weeks after PFF and/or CA074me treatment. C) Number of differentially abundant proteins (Log2-fold change > 0.25, adjusted p value < 0.05, N = 3 replicates) and GO-term analysis of whole-cell proteomics conducted on DA neurons treated for 3-weeks with PFF and/or CA074me. Bonferroni-corrected t-tests, ** p < 0.01, *** p < 0.001, **** p < 0.0001.

**Figure 3:**
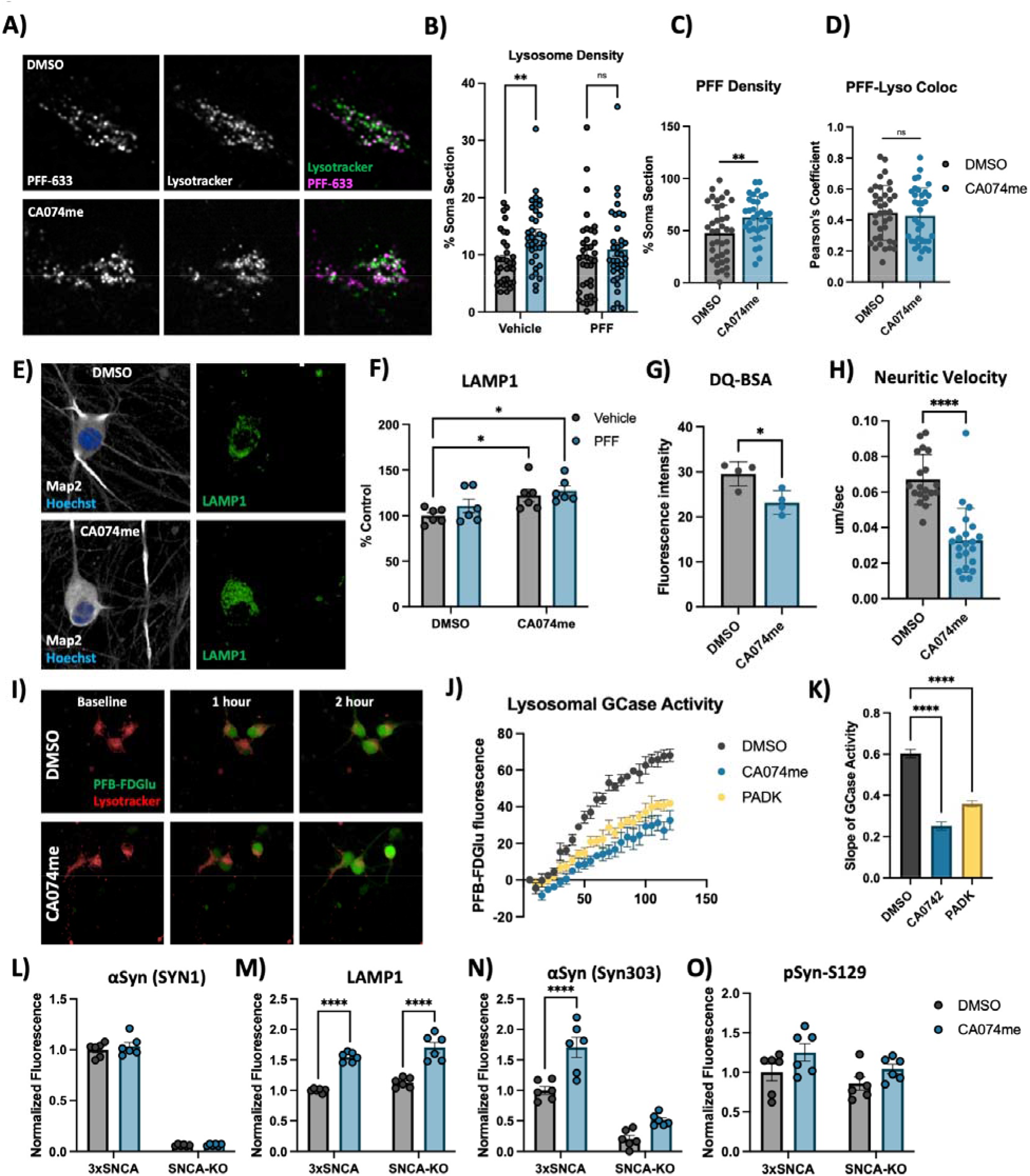
Cathepsin B inhibition increases lysosome abundance but impairs function in dopaminergic neurons. A) Representative live-cell confocal images of neuronal cell bodies stained with lysotracker-green 72-hours after exposure to alexa-633 labelled α-syn PFFs (80 nM). B) Lysosome density per cell body, measured as the percentage of lysotracker-positive area per cell soma. C) PFF density per cell body, measured as the percentage of PFF-633-positive area per cell soma. D) Colocalization of lysotracker and PFF-633 measured using Pearson’s coefficient per cell soma. E) Representative immunofluorescent images from high-content confocal imaging of DA neurons treated with CA074me (1 μM) and/or α-syn PFFs (300 nM) for 3 weeks and stained for Map2 and LAMP1. F) Quantification of LAMP1 in Map2-positive cells. G) Lysosomal degradative capacity measured by fluorescence intensity of DQ-BSA fluorogenic probe 24-hours after dye loading. H) Quantification of lysosome velocity in neurites measured by live-cell confocal imaging and quantified using TrackMate. Points represent individual quantified image fields derived from 6 independent experiments. I) Representative images of neurons stained with lysotracker deep-red and PFB-FDGlu fluorescent signal at baseline and 1– or 2-hours after dye-loading. J) Quantification of PFB-FDGlu fluorescence per cell in DA neurons pre-treated for 24-hours with CA074me or PADK. K) Quantification of the slope of PFB-FDGlu fluorescence versus time. L-O) Immunofluorescent quantifications in α-syn triplication DA neurons versus SNCA-KO control of L) total α-syn levels with SYN1 antibody, M) LAMP1, N) α-syn aggregates with SYN303 antibody, or O) pSyn-S129. T-test or Bonferroni-corrected t-tests, * p < 0.05, ** p < 0.01, **** p < 0.0001.

For PFF treatments on RPE1 cells, 50,000 cells were plated on 12-well plates. After 24 hours, PFF was added and cells were allowed to continue growing for 48 hours before cells were washed with PBS and dissociated with trypsin to remove non-internalized PFFs before being lysed in RIPA buffer.

For PFF high-content imaging assays with PFF, neurons were treated with PFF after 2 weeks of differentiation in 96-well plates. After treatment, media was refreshed every 5-7 days normally. At completion of treatment, cells were washed with PBS and fixed with 4% paraformaldehyde.

### High-content imaging – immunofluorescence

Cells were permeabilized for 10 minutes with 0.3% saponin (lysosome immunostaining) or 0.2% triton X-100 (pS129-α-syn assay and TFEB assay) in PBS and blocked with 1% BSA, 4% goat-serum and 0.02% triton X-100 in PBS. Antibodies used are described in Supplemental Table 7. High content imaging was performed on an Opera Phenix high-content confocal microscope (Perkin Elmer) and image analysis was performed using Columbus (Perkin Elmer). Data processing was then conducted using R studio as previously described ^52^. Briefly, nuclei were first identified by the Hoechst channel, and surrounding soma was identified as Map2-positive region. Relevant secondary stains were then quantified within this Map2-defined region. Single-cell data were then processed using a custom script in R studio to filter objects based on nuclear size, nuclear shape and Map2 staining intensity to identify only the neuronal cells for inclusion in subsequent analysis.

### High content imaging – live cell assays

PFB-FDGlu GCase activity assay: Cells in 96-well plates were pre-loaded for 30 minutes with lysotracker deep red (1:20,000, Invitrogen). Media was then exchanged for FluoroBrite imaging media (Thermo) containing 25uM of PFB-FDGlu (Invitrogen) and cells were then imaged on the Opera Phenix every 15 minutes for 2 hours to monitor GCase activity. Using the Columbus software, lysotracker signal was used to identify cells for quantification of GCase substrate fluorescence, which is depicted as the mean fluorescence per cell.

DQ-BSA: Cells were pre-loaded with DQ-BSA (Invitrogen) for the indicated duration in standard culture media. Cells were then stained with lysotracker deep red (1:20,000, Invitrogen) for 30 minutes and media was exchanged for FluoroBrite and imaging was conducted on the Opera Phenix. Using the Columbus software, lysotracker signal was used to identify cells and DQ-BSA fluorescence intensity was measured.

### Whole-cell proteomics mass spectrometry

For proteomics experiments on iPSC-derived DA neurons, 750,000 neurons were plated on 6-well plates and differentiated for 3 weeks. After 3 weeks, neurons were treated with CA074me and/or 300 nM of PFF and then maintained normally for 3 weeks. No additional drug or PFF were added during the maintenance period. Sample processing, mass-spectrometry and data analysis was performed by The Proteomics and Molecular Analysis Platform at the Research Institute of the McGill University Health Centre (RI-MUHC). Samples were processed for TMT labelling according to the manufacturer recommendations (ThermoFisher TMT-16plex reagents). Labelled peptides were fractionated using Pierce™ High pH Reversed-Phase Peptide Fractionation Kit into 8 fractions. Each fraction was re-solubilized in 0.1% aqueous formic acid and 2 micrograms of each was loaded onto a Thermo Acclaim Pepmap (Thermo, 75uM ID X 2cm C18 3uM beads) precolumn and then onto an Acclaim Pepmap Easyspray (Thermo, 75uM X 15cm with 2uM C18 beads) analytical column separation using a Dionex Ultimate 3000 uHPLC at 250 nl/min with a gradient of 2-35% organic (0.1% formic acid in acetonitrile) over three hours running the default settings for MS3-level SPS TMT quantitation ^53^ on an Orbitrap Fusion instrument (ThermoFisher Scientific) operated in DDA-MS3 mode.

To translate.raw files into protein identifications and TMT reporter ion intensities, Proteome Discoverer 2.2 (ThermoFisher Scientific) was used with the built-in TMT Reporter ion quantification workflows. Default settings were applied, with Trypsin as enzyme specificity. Spectra were matched against the human protein fasta database obtained from Uniprot(2022). Dynamic modifications were set as Oxidation (M), and Acetylation on protein N-termini. Cysteine carbamidomethyl was set as a static modification, together with the TMT tag on both peptide N-termini and K residues. All results were filtered to a 1% FDR.

Pathway analysis of differentially abundant proteins was conducted using STRING ^54^.

### Western Blot

Cultured cells were washed with PBS and collected in RIPA lysis buffer with protease inhibitors. Protein concentration was determined using the Pierce™ BCA Protein Assay Kit (Thermo Scientific™) and proteins were prepared at the desired concentration in 6X Laemmli buffer and heated at 95°C for 5 minutes. 20 μg of protein were loaded on polyacrylamide gels, run with SDS running buffer and transferred onto nitrocellulose membranes and blocked for 30 mins in 5% skim milk made in 1X PBS with 0.1% Tween. For alpha-synuclein blots membranes were fixed using 4% PFA and 0.1% glutaraldehyde for 30 minutes before blocking. Blocked membranes were incubated with primary antibody (Supplemental Table 7) at 4°C overnight followed by HRP-conjugated secondary antibodies for 90 mins at room temperature. Protein detection was performed by chemiluminescence using Clarity Western ECL Substrate (Biorad) and Western blots were quantified using ImageJ.

### RNA Extraction and qRT-PCR

RNA isolation was performed using the RNeasy Mini Kit (Qiagen) and cDNA was generated by RT-PCR using the MMLV Reverse Transcriptase kit (Thermo) with random hexamer primers. Real-time quantitative PCR was performed using SSoAdvanced SYBR Green Master Mix (Biorad). Primers for specified target genes were designed using NCBI PrimerBlast (Supplemental Table 8).

### Live cell confocal imaging and analysis

Neurons were plated on CELLview™ 4-chamber imaging dishes (Greiner) at 100k cells per well. After 3 weeks of differentiation neurons were treated with alexa633 labelled PFF and/or CA074me. After 72 hours neurons were washed and incubated with 50nM of Lysotracker™ Green DND-26 (Invitrogen) for 30m at 37 °C in standard culture media. The dye solution was exchanged for FluoroBrite™ DMEM (Gibco) and plates were immediately imaged. Images were acquired on a custom Andor spinning disc confocal microscope at 100X magnification. Single frames were acquired for cell bodies (488nM Lysotracker, 647 PFF). For neuronal trafficking movies, frames were acquired every 1. 5s for a total of 61 frames.

For the analysis of somatic lysotracker and PFF colocalization cell bodies were masked manually using FIJI ImageJ. For each image, Lysotracker and PFF signal were binarized using Otsu automatic thresholding, and binarized co-cluster signal was obtained using the “Image Calculation>>AND” function. Somatic slice densities of Lysotracker, PFF, or co-clusters were calculated via “Analyze Particles”. Finally, percentage of PFF in lysosomes was obtained by normalizing co-cluster particle density to Lysotracker density, and percentage of lysosomes containing PFF was similarly obtained by normalizing co-cluster particle density to PFF particle density.

Lysosomal motility was analyzed using the FIJI ImageJ TrackMate plugin ^55,56^. The generated tracks were then filtered by max track speed and then analyzed using Python.

### Electron Microscopy

RPE1 cells grown in Lab-Tek chambers (Nunc) were rinsed in 0.1M Na Cacodylate buffer and fixed with 2.5% glutaraldehyde in 0.1M Na Cacodylate for 24 hours at 4°C. Cells were then post-fixed with 1% aqueous osmium tetroxide (Mecalab) for 1 h at 4°C, and stained with 4% uranyl acetate (EMS) in 70% ethanol for 45min at 4°C. After dehydrations in ascending alcohols, cells were embedded in Epon resin (Mecalab), and cut at 75 nm thickness in the ultra microtome. Sections were collected in 200 Mesh cooper grids (EMS) and stained with 4% uranyl acetate for 5min following by Reynold’s lead citrate for 2 min. Sections were visualized using a transmission electron microscope (Tecnai G2 Spirit Twin 120 kV Cryo-TEM) coupled to a camera (Gatan Ultrascan 4000 4 k × 4 k CCD Camera model 895). The identification of cellular elements was based on standard descriptions ^57^.

### Statistical Analysis

Statistical analysis was conducted in GraphPad Prism9 software. For experiments with iPSC-derived neurons biological replicates were defined as experiments conducted at different times from the same batch of banked NPCs, or as experiments conducted in parallel from different batches of NPCs. A minimum of 3 distinct batches of NPCs were used for each experiment.

Statistical comparisons were performed using t-tests (only 2 conditions), Bonferroni-corrected t-tests (more than 2 conditions compared). Significance levels are depicted in figure legends.

## Results

### Variants in *CTSB* likely drive the association with PD and are associated with *CTSB* expression in multiple brain regions

Variants in the genetic locus containing *CTSB* are significantly associated with risk of PD ^9^ yet this locus includes multiple other genes including *FDFT1*, *NEIL2*, *GATA4* and it remains uncertain whether *CTSB* itself drives the association. We examined all the variants that are 1MB upstream or downstream to the top GWAS variant in this locus and using GCTA-COJO, we show that an intronic *CTSB* variant (rs1293298, *p*=3.41E-16, located in intron 1 of *CTSB* within a potential enhancer region) is the top variant associated with PD risk, without secondary associations. Fine mapping using FINEMAP gave this variant the highest posterior probability (0.127) of being causal, of all nominated variants. This variant is in LD with multiple variants in this locus (r^2^>0.8) that are associated with *CTSB* expression with H4 posterior probability >0.8 in multiple brain regions. The associations between genetic variants, PD and *CTSB* expression in PD-relevant brain regions such as basal ganglia, cortex and nucleus accumbens are depicted in Figure 1. In particular, the minor allele of the rs1293298 *CTSB* variant linked to PD in GWAS exhibits a protective effect against PD and is associated with elevated expression levels of *CTSB* in brain tissues relevant to the disease (Fig 1A-D). Analysis using SMR and HEIDI suggests that the QTLs in *CTSB* are potentially causally linked to PD with p HEIDI > 0.05 in multiple tissues (i.e., we could not reject the null hypothesis that there is a single causal variant affecting both gene expression and risk for PD). All results from the GCTA-COLOC, FINEMAP, SMR and HEIDI analyses are detailed in Supplementary Tables 9-11.

These common *CTSB* variants occur in non-coding regions and likely exert their effects through altering expression levels. However, given the evidence that protective *CTSB* variants are associated with increased mRNA expression levels, we hypothesized that loss of function coding variants in *CTSB* would be likely to promote PD risk. We conducted rare variant analysis in 5,801 PD cases and 20,427 controls from six cohorts (Supplementary Table 2). We observed a nominal association between all rare variants and variants with high CADD score and PD risk in the Sheba cohort (p=0.03 and p=0.049, respectively). However, upon examining other cohorts and conducting a meta-analysis we did not find any additional associations (Supplemental Table 12). Additionally, we studied the role of rare *CTSB* variants on the age of PD onset. We found nominal association between functional variants and age at onset in McGill cohort (p=0.044) and in the meta-analysis for functional and non-synonymous variants (p=0.048 and p=0.043, respectively). All these results should be interpreted with caution as no p-values survived multiple comparisons.

### CatB inhibition promotes **α**-syn aggregation in dopaminergic neurons

To functionally interrogate the role of catB in the handling of α-syn fibrils we generated iPSC-derived dopaminergic neurons ^41,42^ and treated them with pre-formed α-syn fibrils (PFFs) and the catB inhibitor CA074me (1 μM) ^58^. Exposure to PFFs promotes endogenous α-syn aggregation which can recapitulate many features of Lewy pathology including the accumulation of S129-phosphorylated α-syn (pSyn-S129) ^59^. We used high-content confocal imaging to quantify pSyn-S129 in Map2-positive neurons following exposure to PFF and/or CA074me (Fig 2A). After 2, 3 or 4 weeks, a single treatment with CA074me administered at the time of PFF exposure increased the abundance of pSyn-S129 (Fig 2B). Similar effects were observed with PADK, a distinct catB inhibitor (Fig S4A). CA074me did not affect total α-syn levels (Fig S4B) and no significant loss of TH+ dopaminergic neurons was observed 4 weeks post-treatment (Fig S4C). The induction of pSyn-S129 was abolished in neurons lacking endogenous α-syn (SNCA-KO) (Fig S4D), indicating that α-syn seeding, rather than phosphorylation of PFFs themselves gave rise to this pSyn-S129 signal.

We performed whole-cell proteomics with TMT labelling to characterize the broader impact of PFF or CA074me treatment on human DA neurons (Supplementary Table 13). We found that 3 weeks after treatment CA074me had minimal residual effects, while PFF exposure significantly altered the abundance of 60 proteins (Fig 1C – Venn diagrams). GO-term analysis with STRING^54^ revealed that the predominant pathways impacted by PFF treatment were the downregulation of proteins involved in cellular adhesion and cytoskeletal organization (Fig 1C – bar graphs). Combining CA074me with PFF resulted in >2× the number of differentially abundant proteins than either treatment alone. GO-term enrichment revealed many similar pathways were downregulated by either PFF or PFF+CA074me, but the combined treatment also upregulated pathways not found to be altered by PFF alone.

### CatB inhibition induces lysosome dysfunction in dopaminergic neurons

Extracellular α-syn aggregates are taken into neurons by a variety of mechanisms and are rapidly trafficked to lysosomes ^60,61^. To determine whether CA074me treatment altered the trafficking of PFFs into lysosomes or their persistence there, we performed live cell confocal imaging of DA neurons exposed to alexa-633 tagged PFFs (PFF-633) for 72 hours and stained with lysotracker (Fig 3A). CA074me increased the overall lysosome content (Fig 3B) and the density of PFF-633 fluorescent puncta per cell (Fig 3C) but colocalization of PFF-633 with lysosomes was unchanged (Fig 3D). We interpret these observations as indicating that although the abundance of both lysosomes and PFF-633 within each cell is slightly elevated in CA074me treated neurons, the proportion of PFF-633 trafficked to lysosomes is unaffected.

Given the observed increase in lysotracker density, we next sought to further characterize how catB inhibition affected lysosome abundance and function in human DA neurons. Similar to lysotracker, the abundance of the lysosomal membrane protein LAMP1 was increased after CA074me treatment, independent of concurrent PFF exposure (Fig 3E, F). However, the degradative capacity of lysosomes (measured using the fluorogenic probe DQ-BSA) was reduced (Fig 3G). The speed of lysosomal trafficking in neuritic projections was also reduced following CA074me (Fig 3H). Lastly, given the genetic interaction between variants in *CTSB* and *GBA1* in PD risk ^10^ and that catB has been found to regulate glucocerebrosidase (GCase) activity in HEK293 cells ^62^ we assessed the impact of catB inhibition on lysosomal GCase activity in DA neurons using the fluorogenic probe PFB-FDGlu (Fig 3I-K). CatB inhibition with either CA074me or PADK impaired lysosomal GCase activity (Fig 3J, K). These observations indicate that despite increasing lysosome abundance, catB inhibition impairs several aspects of lysosome function in DA neurons, including degradative capacity, trafficking and GCase activity.

To determine whether altered lysosome function could be related to accumulation of α-syn aggregates, (which has been found to impact lysosomal hydrolase trafficking ^11,13^) we differentiated 3×SNCA and SNCA-KO iPSCs ^45^ and treated them with CA074me for 3 weeks. We observed that while total levels of α-syn were unchanged by CA074me (Fig 3L), LAMP1 was increased in both 3×SNCA and SNCA-KO neurons (Fig 3M) indicating the increase in lysosome content is independent of α-syn. We additionally stained with antibodies that preferentially detect aggregated species of α-syn (Syn303) and found this to be increased in CA074me-treated SNCA-triplication neurons (Fig 3N), although almost no S129-pSyn signal above background was detected in these cells (Fig 3O).

### CTSB levels regulate PFF clearance in RPE1 cells

While CA074me is selective for catB, it has been reported to inhibit other cathepsins, albeit at concentrations greater than those used in this study ^63,64^. RNA sequencing of our iPSC-derived DA neurons indicated that 12 cathepsin family members were expressed at the RNA level (data not shown), including *CTSD* and *CTSL* which were previously found to cleave α-syn *in vitro* ^18–20^. To determine how individual cathepsin species contribute to fibrillar α-syn clearance in a cellular context we used CRISPR-interference (CRISPRi) to generate RPE1 cell lines in which *CTSB*, *CTSD*, *CTSL* or α-syn (*SNCA*) were stably repressed (denoted CTSBi, CTSDi, CTSLi and SYNi respectively) as well CRISPR-activation (CRISPRa) to upregulate *CTSB* (CTSBa) (Fig 4A,B). Endogenous α-syn protein was undetectable in SNCAi cells and modestly elevated in CTSBi cells (Fig 4C, D). In contrast, CTSBa had no effect on endogenous α-syn protein (Fig 4E, F). Strikingly, 48 hours after exposure to PFFs, CTSBi cells (but not CTSDi or CTSLi) exhibited significantly greater accumulation of α-syn aggregates compared to control cells (Fig 4G,H) while CTSBa had the opposite effect, modestly reducing the levels of aggregated α-syn (Fig 4I, J). This effect was recapitulated by treatment of control or SNCAi cells with CA074me (Fig S5A,B) indicating that this accumulation reflects either increased cellular uptake or failed clearance of the PFFs, rather than new aggregate seeding.

**Figure 4:**
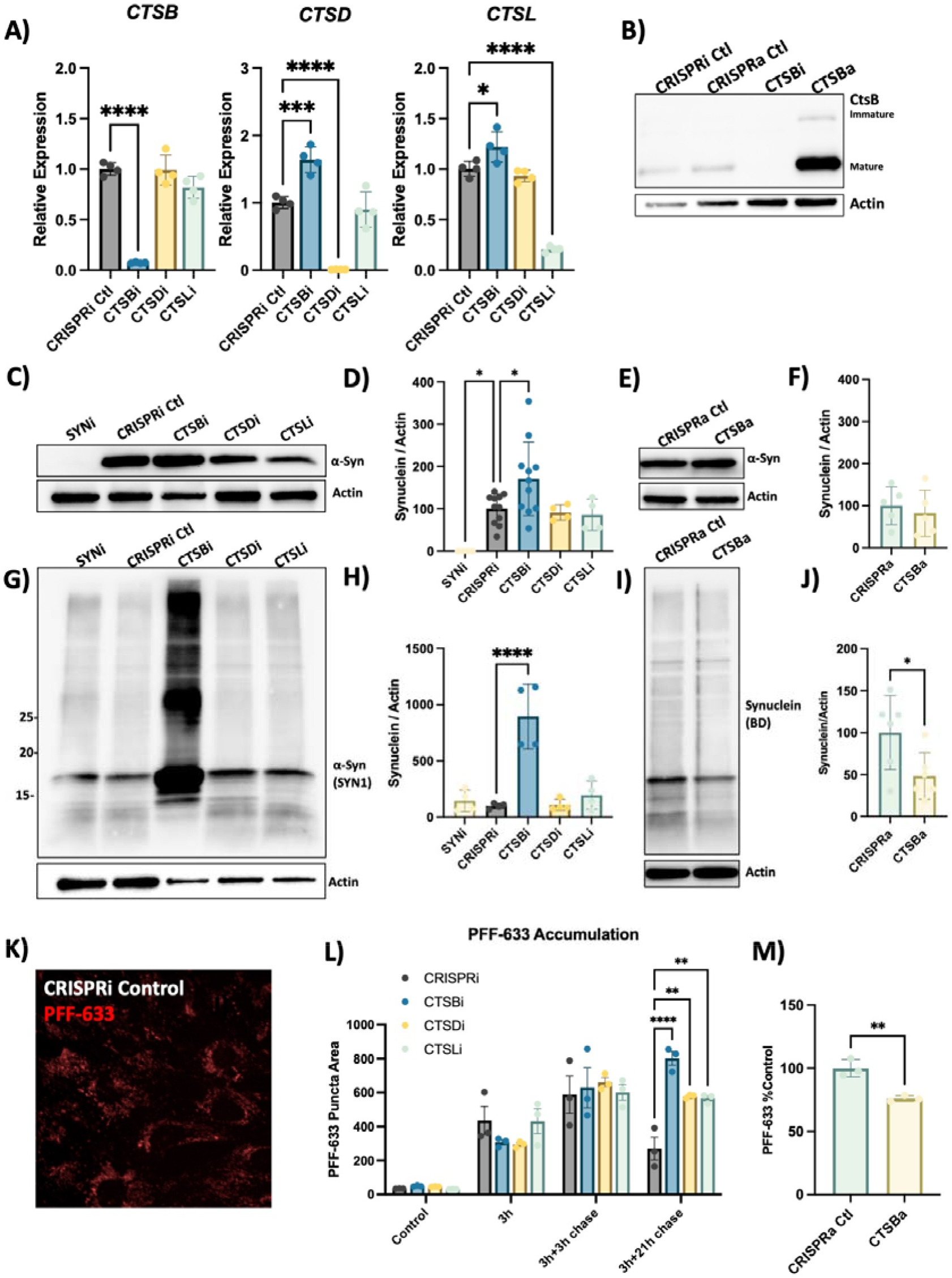
CTSB but not *CTSD* or *CTSL* repression impairs PFF clearance in RPE1 cells. A) Validation of target knockdown in CRISPRi RPE1 cells. *CTSB*, *CTSD* and *CTSL* mRNA levels were measured by qPCR in control (CRISPRi Ctl), *CTSB*-knockdown (CTSBi), *CTSD*-knockdown (CTSDi) and *CTSL*-knockdown (CTSLi) cell lines. B) Western blots depicting protein levels of cathepsin B (catB) in CRISPRi control, CRISPRa control, *CTSB* knockdown (CTSBi) and *CTSB* upregulation (CTSBa) RPE1 cell lines. C) Representative western blot depicting protein levels of α-syn (SYN1 antibody) and actin. D) Western blot quantification depicting protein levels of α-syn relative to actin expressed as percentage of CRISPRi control. E) Representative western blot depicting protein levels of α-syn and actin. F) Western blot quantification depicting protein levels of α-syn relative to actin expressed as percentage of CRISPRa control. G) Representative western blot depicting protein levels of α-syn (SYN1 antibody) and actin 48-hours after treatment of RPE1 cell lines with 300nM of α-syn PFFs. H) Western blot quantifications depicting levels of α-syn (SYN1 antibody – quantification of whole lane) relative to actin in PFF treated RPE1 cells. I) Representative western blot depicting protein levels of α-syn (SYN1 antibody) and actin 48-hours after treatment of RPE1 cell lines with 300nM of α-syn PFFs. J) Western blot quantifications depicting levels of α-syn (SYN1 antibody – quantification of whole lane) relative to actin in PFF treated RPE1 cells. K) Representative image of CRISPRi control RPE1 cells 48-hours after treatment with alexa-633 tagged α-syn PFFs (80 nM). L,M) Quantification of PFF-633 fluorescent intensity per cell in RPE1 cell lines. T-test or Bonferroni-corrected t-tests, * p < 0.05, ** p < 0.01, *** p < 0.001, **** p < 0.0001.

To determine whether the uptake or clearance of PFFs was affected, we used Alexa-633 fluorescently labelled PFFs and conducted a pulse-chase experiment to monitor the uptake and subsequent clearance of PFFs (Fig 4K, L). During a 3h exposure, or 3-hour exposure with short washout (3h chase), the PFF-633 levels per cell were similar across cell lines. However, when washout was extended to 21-hours CTSBi cells retained more PFF-633 (Fig 4L), suggesting impaired clearance. CTSDi and CTSLi also appeared to impair PFF clearance in this assay but to a lesser extent than CTSBi. Similar to what we observed by Western blot, when we exposed CTSBa cells to PFF-633 for 48 hours, we observed a reduced accumulation of the tagged PFF (Fig 4M). Taken together, these findings indicate that CatB regulates the clearance of internalized α-syn aggregates in lysosomes.

### CTSB repression impairs autophagy and lysosomal function in RPE1 cells

We next used our CTSBi cell line to further interrogate the role of CTSB in regulating lysosome abundance and overall lysosome function. Given the accumulation of lysosomal structures in DA neurons we hypothesized that loss of catB caused a state of lysosomal dysfunction that resulted in the accumulation of hypo-functional lysosomes. However, a previous study found that loss of catB also triggered lysosome biogenesis, and in select circumstances had a net effect of enhancing clearance of lysosomal cargo ^65^. CTSBi cells had significant accumulation of lysosomes, as indicated by an increase in LAMP1 and LAMP2 immunofluorescent signal (Fig 5A-C), an increase in LAMP1 and GCase protein levels (Fig 5D-F) and increased number and size of electron dense lysosome-like structures (including lysosomes and multivesicular bodies) observed by electron microscopy (Fig 5G). However, despite having increased lysosome content and increased GCase protein levels, the activity of lysosomal GCase per cell was significantly reduced in CTSBi cells (Fig 5H, I).

**Figure 5:**
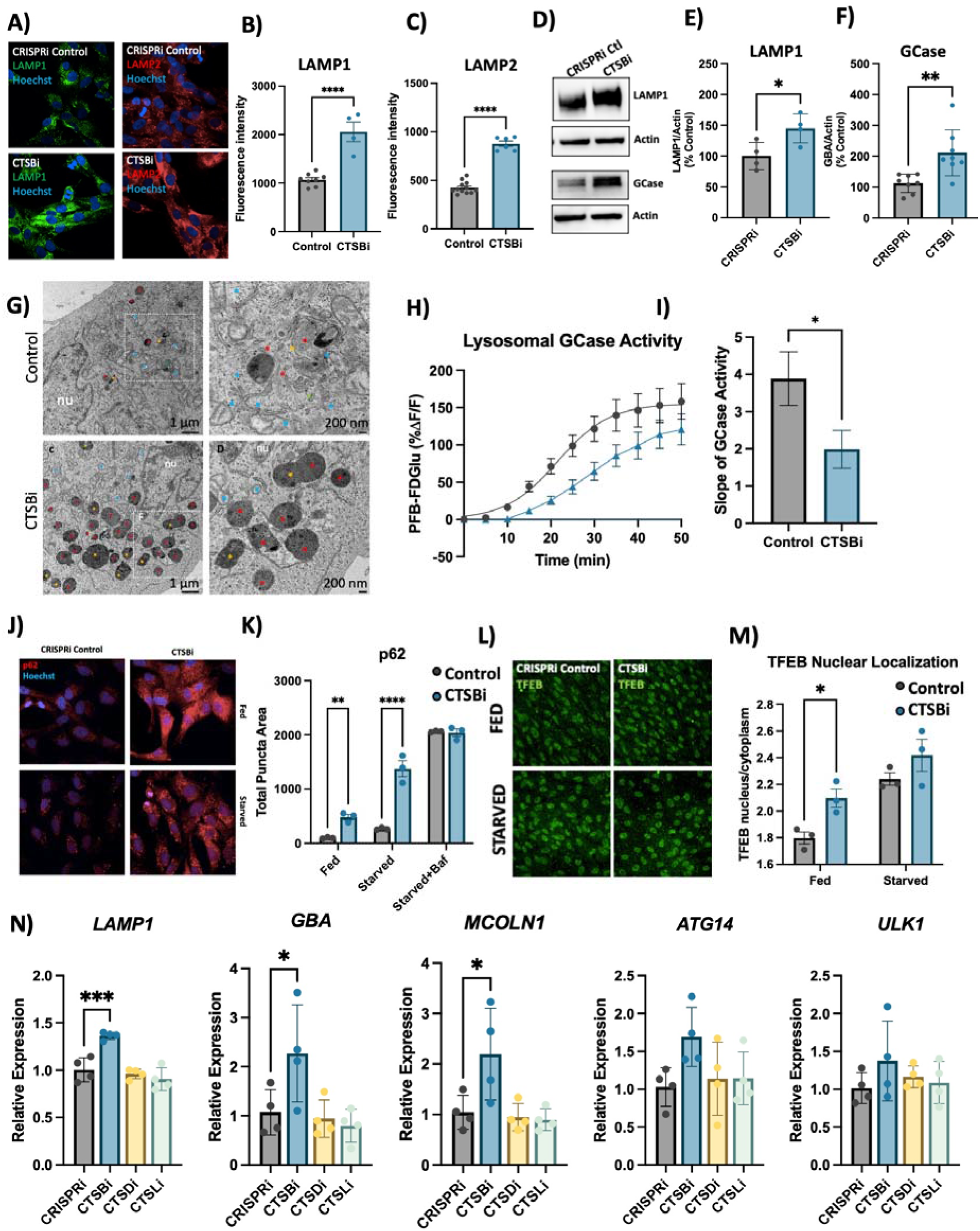
CTSB repression increases lysosome abundance but impairs function in RPE1 cells. A-C) Representative images and quantifications of LAMP1 and LAMP2 immunofluorescence in CRISPRi control and CTSBi RPE1 cells. D-F) Representative western blots and quantifications depicting protein levels of LAMP1 and GCase/*GBA* relative to actin. G) Electron microscopy images of CRISPRi and CTSBi RPE1 cells. Electron-dense multivesicular bodies-like structures are indicated by red asterisk, lysosomes by orange asterisk, secondary lysosomes by green asterisk, whereas mitochondria are indicated by blue asterisk. H) GCase activity measured as PFB-FDGlu fluorescence intensity over time. I) Quantification of the slope of the PFB-FDGlu fluorescence versus time curves. J, K) Representative immunofluorescent images and quantification of p62 area per cell in RPE1 cell lines (CRISPRi control – grey bars, and CTSBi – blue bars) under fed, 16-hour starvation and 16-hour starvation with bafilomycin conditions. L, M) Representative images of TFEB immunofluorescence and quantification of nuclear TFEB (overlapping with Hoechst nuclear stain) relative to cytoplasmic TFEB per cell under fed and starved conditions. N) RNA expression levels of the indicated genes in RPE1 cell lines measured by RT-qPCR expressed relative to CRISPRi Control. Bonferroni-corrected t-tests, * p < 0.05, *** p < 0.001. T-test or Bonferroni-corrected t-tests, * p < 0.05, ** p < 0.01, **** p < 0.0001.

To determine whether impairment in autophagic flux could contribute to the accumulation of lysosomes, we measured the abundance of p62 puncta under fed and starved conditions, and in the presence of bafilomycin (to inhibit lysosomal clearance of autophagosomes). CTSBi resulted in increased abundance of p62 puncta under fed and serum-starved conditions, but not in the presence of bafilomycin (Fig 5J, K) suggesting an impairment in the clearance of autophagosomes. Similarly, we observed accumulation of the autophagy-associated proteins LC3B and p62 by western blot in CTSBi cells (but not CTSDi or CTSLi), in the absence of serum starvation (Fig S6A,B).

While this impaired autophagosome clearance likely contributes to the accumulation of lysosomes in CTSBi cells, catB has previously been reported to regulate lysosome biogenesis via activation of the transcription factor TFEB ^65,66^. Given that mRNA levels of *CTSD* and *CTSL* were already noted to be increased in CTSBi cells (Fig 4A) we suspected a role for increased TFEB activity. Indeed, we found that in CTSBi cells, nuclear localization of TFEB was increased in non-starved cells (Fig 5L, M) and mRNA levels of several lysosomal genes (including *LAMP1*, *GBA* and *MCOLN1*) were transcriptionally upregulated (Fig 5N). These results taken together indicate that a combination of impaired lysosome turnover and upregulated lysosome biogenesis contribute to the increased abundance of dysfunctional lysosomes in RPE1 cells lacking *CTSB*.

### Knockout of CTSB in human dopaminergic neurons leads to lysosomal dysfunction

To further confirm the lysosomal phenotypes that we previously observed in neurons treated with CA074me, we generated *CTSB* knockout iPSCs (CTSB-KO) and differentiated them into dopaminergic neurons (Fig 6A, B). Similar to CA074me treatment, in CTSB-KO neurons lysosome abundance measured either by LAMP1 immunofluorescence (Fig 6C) or lysotracker (Fig 6D) were increased, while degradative capacity (Fig 6E) and neuritic trafficking velocity (Fig 6F) were reduced. Lysosomal GCase activity was likewise reduced by approximately 20% in CTSB-KO neurons (Fig 6G, H) although GCase protein levels were unaffected (Fig 6I, J).

**Figure 6:**
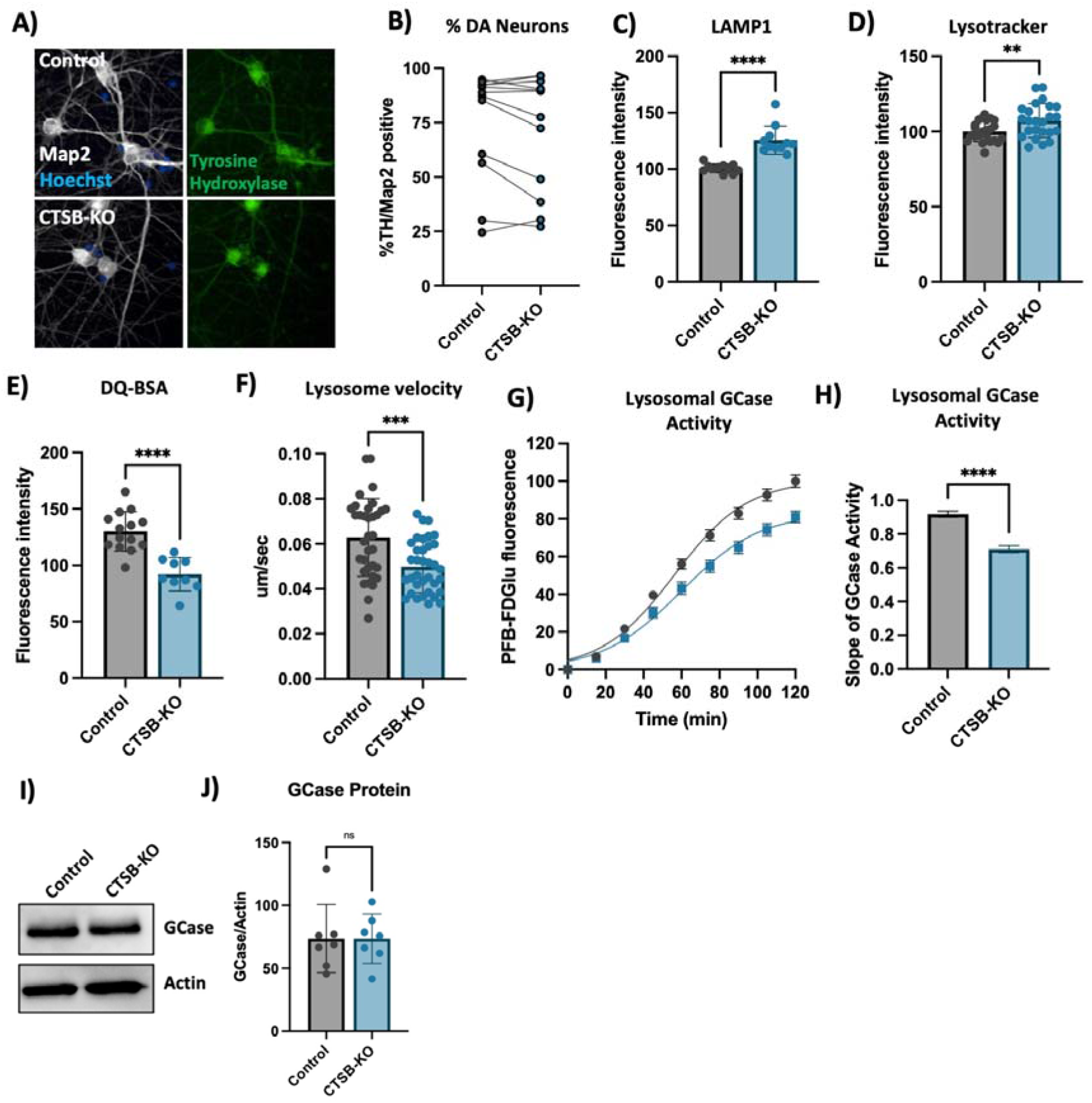
CTSB knockout impairs lysosome function in dopaminergic neurons. A) Representative immunofluorescent images from high-content confocal imaging of DA neurons differentiated from control or CTSB-KO iPSCs and stained for Map2, tyrosine hydroxylase (TH) and α-syn. B) Quantification of the percentage of TH and Map2-positive cells in matched batches of iPSC-derived neurons. C) High-content imaging-based quantification of LAMP1 immunofluorescence per cell in Map2-positive iPSC-derived DA neurons. D) Quantification of lysotracker fluorescence in iPSC-derived DA neurons. E) Lysosomal degradative capacity measured by fluorescence intensity of DQ-BSA fluorogenic probe 24-hours after dye loading. F) Quantification of lysosome velocity in neurites measured by live-cell confocal imaging and quantified using TrackMate. Points represent individual quantified image fields derived from 6 independent experiments. G) Quantification of PFB-FDGlu fluorescence per cell in iPSC-derived DA neurons H) Quantification of the slope of PFB-FDGlu fluorescence versus time. I, J) Representative western blot and quantification of GCase and actin in iPSC-derived DA neurons. T-tests, ** p < 0.01, *** p < 0.001, **** p < 0.0001.

Unlike CTSB-knockdown RPE1 cells, CTSB-KO neurons did not exhibit detectable activation of TFEB (Fig S7A) or transcriptional upregulation of lysosomal genes (Fig S7B) arguing that catB may regulate TFEB activity and lysosome biogenesis in a cell-type or context-dependent manner.

### CTSB deficiency promotes synuclein pathology in human dopaminergic neurons and midbrain organoids

CTSB-KO DA neurons were found to have modestly elevated levels of endogenous α-syn (Fig 7A,B). When treated with PFFs for 72 hours CTSB-KO neurons did not exhibit higher levels of total α-syn (Fig 7A, C). However, 3 and 4 weeks after PFF treatment CTSB-KO neurons accumulated significantly more pSyn-S129, and this was evident when measuring either the average pSyn-S129 intensity within Map2-positive neurons, or the percentage of pSyn-positive cell bodies (Fig 7D-F). The efficiency of PFF uptake (measured by alexa-488 tagged PFF internalization) was unaffected in CTSB-KO neurons (Fig 7G,H) but as expected LAMP1 was elevated (Fig 7G,I). Using live-cell confocal microscopy, we observed a modest increase in total PFF-633 levels in CTSB-KO neurons 72 hours after treatment (Fig S8A, B), however there was no difference in the trafficking of alexa-633 tagged PFFs to lysosomes as measured by PFF-633 and lysotracker colocalization (Fig S8C).

**Figure 7:**
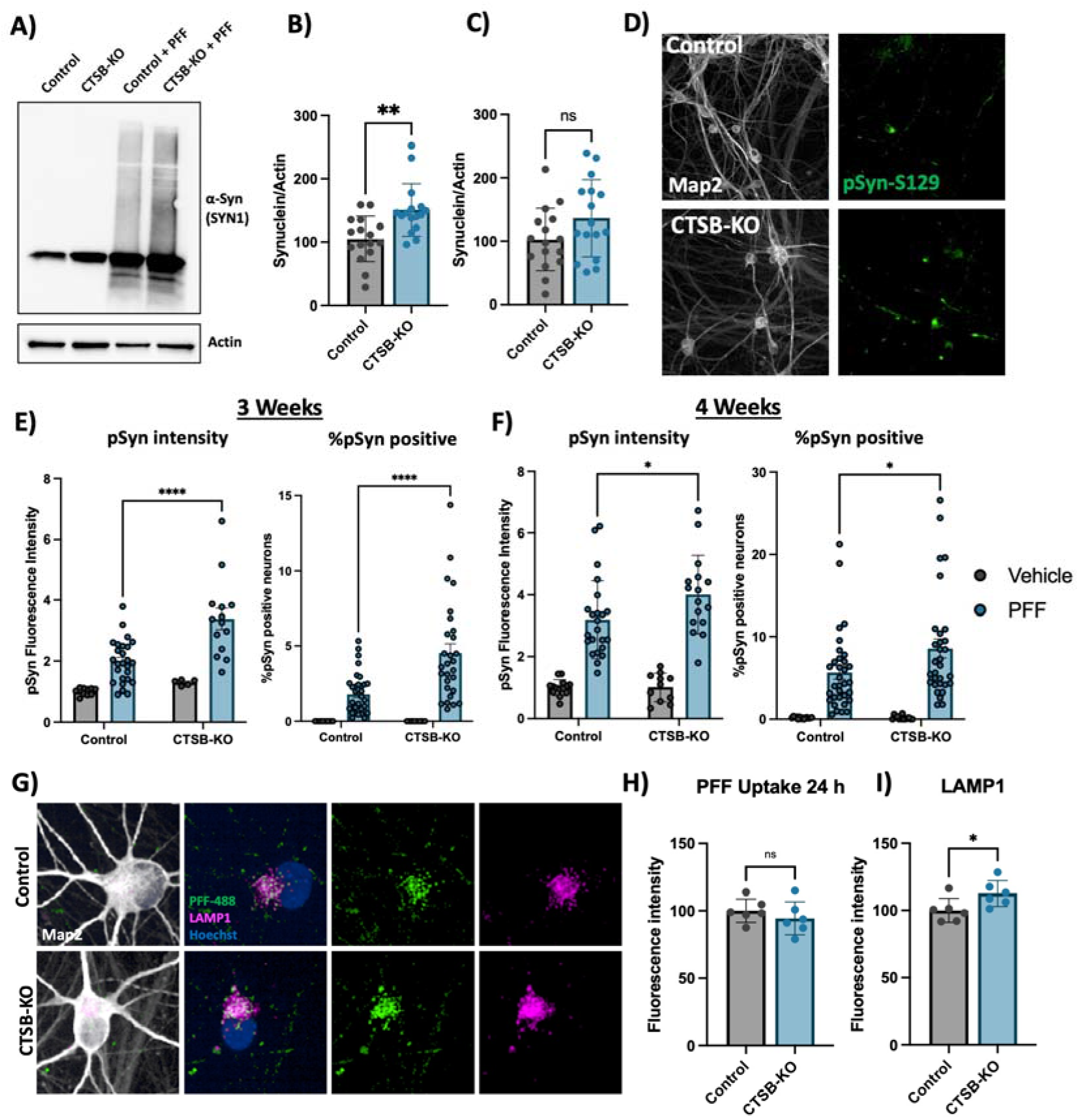
CTSB knockout enhances the effect of α-syn PFFs on dopaminergic neurons. A) Representative western blot showing levels of α-syn (SYN1 antibody) in untreated control or CTSB-KO DA neurons as well as PFF treated DA neurons. B, C) Western blot quantifications depicting protein levels of α-syn relative to actin for endogenous α-syn (B) or PFF (C), normalized to the respective Control. D) Representative immunofluorescent images from high-content confocal imaging of Control or CTSB-KO DA neurons treated with α-syn PFFs (300 nM) and stained for Map2 and pSyn-S129. E, F) Quantification of pSyn-S129 in Map2-positive cells 3-weeks (E) and 4-weeks (F) after PFF treatment. Left graphs depict pSyn-S129 fluorescence intensity within Map2-positive cells, and right graphs depict the percentage of Map2-positive cell bodies positive for pSyn-S129 aggregates. G) Representative immunofluorescent images from high-content confocal imaging of DA neurons treated with alexa-488 tagged PFFs (PFF-488, 80 nM) for 24 h and stained for LAMP1 and Map2. H) Quantification of PFF-488 fluorescence per Map2-positive cell. I) Quantification of LAMP1 fluorescence per Map2-positive cell. T-tests or Bonferroni-corrected t-tests, ** p < 0.01, **** p < 0.0001.

To determine whether the loss of catB function could promote α-syn aggregation independent of PFF exposure we generated midbrain organoids from patient-derived iPSCs harboring an SNCA triplication mutation (Fig S3). We have previously found that these organoids spontaneously develop pSyn-S129-positive α-syn aggregates after sustained culture ^45^. We treated 3×-SNCA or isogenic SNCA-KO organoids for 60 days with DMSO (vehicle) or 1 μM CA074me and observed an increase in the abundance of pSyn-S129 (measured as the area positive for pSyn-S129 relative to total organoid area) (Fig 8A-C), while treatment had no effect on total neuron content (Map2-positive area) (Fig 8D). These findings further indicate that catB contributes to lysosomal clearance of both endogenous α-syn and exogenously applied α-syn PFFs.

**Figure 8:**
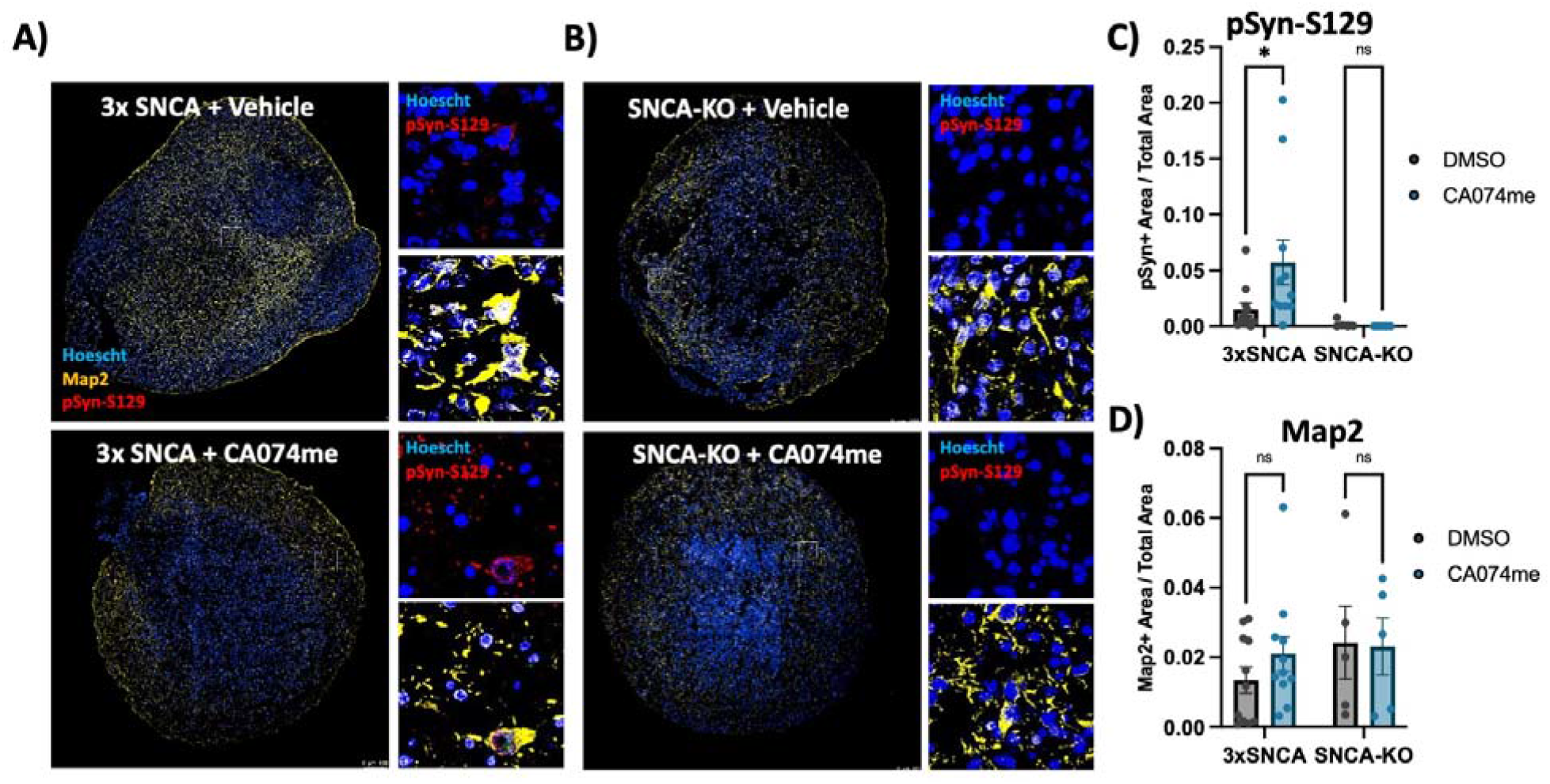
CTSB inhibition promotes pSyn-S129 accumulation in patient-derived midbrain organoids. A, B) Representative immunofluorescent images of Map2 and pSyn-S129 in 5-month old SNCA-triplication (A) and isogenic SNCA-knockout (B) midbrain organoids treated with vehicle (DMSO) or 1 μM CA074me for 60 days. Large images depict representative whole-organoids and high-magnification images depict individual Map2-positive cells. C) Quantification of the pSyn-S129 positive area of the organoid relative to the total organoid size. D) Quantification of the Map2-positive area relative to organoid size. Bonferroni-corrected t-tests, * p < 0.05.

## Discussion

Cathepsin B has previously been suggested to contribute to the degradation of α-syn and genetic variants in the *CTSB* locus are significantly associated with PD, suggesting that this lysosomal protease may play an important role in the disease. Here we provide genetic and functional evidence supporting the crucial involvement of *CTSB* in PD, specifically relating to the function of lysosomes and degradation of α-syn aggregates in dopaminergic neurons.

Firstly, our genetic analysis provides compelling evidence for a causal relationship between common non-coding variants in the *CTSB* gene and both brain expression levels of *CTSB* and PD risk. This genetic analysis indicates that of the variants and genes in the *CTSB* GWAS locus, the association is most likely driven by *CTSB* variants that affect its expression in different brain regions. In particular, the minor allele of the rs1293298 *CTSB* variant, linked to PD in GWAS, exhibits a protective effect against PD and is associated with elevated expression levels of *CTSB* in several brain tissues. This finding is also supported by recent work in which we have used machine learning to nominate the most likely causative genes in each known PD locus, in which *CTSB* was also found to be the top nominated gene ^67^.

One potential mechanism by which *CTSB* variants may influence PD risk is through the ability of catB to cleave and degrade α-syn ^18–20^. However, while several cathepsins appear capable of cleaving α-syn *in vitro, CTSB* alone stand out as a genetic risk factor for PD. By using genetic tools to modulate the expression levels of *CTSB*, *CTSD* and *CTSL* in RPE1 cells, we show that in a cellular context, *CTSB* is particularly critical for the maintenance of lysosome function and clearance of fibrillar α-syn. This is supported by recent the finding that while many cathepsins exhibit redundancy, the sites within α-syn cleaved by *CTSB* are relatively unique, and unlikely to be compensated for by other cathepsins ^20^. This lack of redundancy may explain why *CTSB* stands out as a genetic risk factor and an essential mediator of α-syn clearance.

In addition to a potential direct role of catB in degrading α-syn aggregates, we have also observed that catB impairment leads to lysosome accumulation and broad impairment of lysosome functions, including impaired GCase activity. Variants in *CTSB* and *GBA* interact to mediate genetic risk for PD ^10^ and given the established importance of GCase in mediating risk of synucleinopathy (reviewed in ^68^) this raises the question of whether the impaired α-syn clearance observed following catB impairment is partially mediated by secondary GCase impairment. This loss of GCase activity occurs despite an increase in overall lysosome content, and in the case of RPE1 cells, an increase in GCase protein levels. One potential mechanism linking catB to GCase activity is via the ability of catB to cleave pro-saposin into saposin C which acts as a co-activator of GCase ^62^. Future studies will be required to determine the importance of GCase as a mediator of catB –dependent α-syn clearance, as well as the mechanism of interaction between these lysosomal proteins.

In the present work we have demonstrated using several cellular models that loss of *CTSB* impairs GCase activity and promotes the accumulation or aggregation of α-syn after exposure to preformed α-syn fibrils. These findings complement genetic evidence that *CTSB* variants associated with increased expression levels are protective against the disease and provides potential mechanistic support for the genetic interaction between *CTSB* and *GBA*. Beyond the direct genetic association, impaired catB expression or activity have also been reported in cellular or animal models associated with PD-risk factors like α-syn/*SNCA*, *GBA*, *TMEM175* and *LRRK2* ^11–17,69^. Together this evidence highlights *CTSB* as an important player in the etiology of synucleinopathies such as Parkinson’s disease, and further study of its biology may help to uncover novel therapeutic approaches to this disease.

## Abbreviations

α-syn: alpha-synuclein
CRISPRa: CRISPR-activation
CRISPRi: CRISPR-inhibition
CTSB/catB: cathepsin B
CTSD: cathepsin D
CTSL: cathepsin L
DA: dopaminergic
eQTL: expression quantitative trait loci
GBA1: glucocerebrosidase
GWAS: genome wide association study
iPSC: induced pluripotent stem cell
LAMP: lysososme associated membrane protein
LD: linkage disequilibrium
PD: Parkinson’s disease
PFF: preformed alpha-synuclein fibril
Map2: microtubule associated protein 2
NPC: neuronal progenitor cell
TFEB: transcription factor EB
TH: tyrosine hydrozylase

## Acknowledgements

JJT is supported by a Parkinson Canada postdoctoral research fellowship. KS is supported by Parkinson Canada Movement Disorders clinical fellowship. ZGO is supported by the Fonds de recherche du Québec – Santé (FRQS) Chercheurs-boursiers award and is a William Dawson Scholar. We would also like to thank all members of the Fon and Gan-Or labs as well as the Neuro EDDU for their support and scientific discussions that helped to shape this work.

The access to part of the participants for this research has been made possible thanks to the Quebec Parkinson’s Network (http://rpq-qpn.ca/en/). We would like to thank the participants in the different cohorts for contributing to this study. This research used the NeuroHub infrastructure and was undertaken thanks in part to funding from the Canada First Research Excellence Fund, awarded through the Healthy Brains, Healthy Lives initiative at McGill University, Calcul Québec and Compute Canada. This research has been conducted using the UK Biobank Resource under Application Number 45551. Data used in the preparation of this article were obtained from the Accelerating Medicine Partnership® (AMP®) Parkinson’s Disease (AMP PD) Knowledge Platform. For up-to-date information on the study, visit https://www.amp-pd.org. The AMP® PD program is a public-private partnership managed by the Foundation for the National Institutes of Health and funded by the National Institute of Neurological Disorders and Stroke (NINDS) in partnership with the Aligning Science Across Parkinson’s (ASAP) initiative; Celgene Corporation, a subsidiary of Bristol-Myers Squibb Company; GlaxoSmithKline plc (GSK); The Michael J. Fox Foundation for Parkinson’s Research; Pfizer Inc.; Sanofi US Services Inc.; and Verily Life Sciences. ACCELERATING MEDICINES PARTNERSHIP and AMP are registered service marks of the U.S. Department of Health and Human Services. Genetic data used in preparation of this article were obtained from the Fox Investigation for New Discovery of Biomarkers (BioFIND), the Harvard Biomarker Study (HBS), the Parkinson’s Progression Markers Initiative (PPMI), the Parkinson’s Disease Biomarkers Program (PDBP), the International LBD Genomics Consortium (iLBDGC), and the STEADY-PD III Investigators. BioFIND is sponsored by The Michael J. Fox Foundation for Parkinson’s Research (MJFF) with support from the National Institute for Neurological Disorders and Stroke (NINDS). The BioFIND Investigators have not participated in reviewing the data analysis or content of the manuscript. For up-to-date information on the study, visit michaeljfox.org/news/biofind. The Harvard Biomarker Study (HBS) is a collaboration of HBS investigators [full list of HBS investigators found at https://www.bwhparkinsoncenter.org/biobank/ and funded through philanthropy and NIH and Non-NIH funding sources. The HBS Investigators have not participated in reviewing the data analysis or content of the manuscript. PPMI is sponsored by The Michael J. Fox Foundation for Parkinson’s Research and supported by a consortium of scientific partners: [list the full names of all of the PPMI funding partners found at https://www.ppmi-info.org/about-ppmi/who-we-are/study-sponsors]. The PPMI investigators have not participated in reviewing the data analysis or content of the manuscript. For up-to-date information on the study, visit www.ppmi-info.org. The Parkinson’s Disease Biomarker Program (PDBP) consortium is supported by the National Institute of Neurological Disorders and Stroke (NINDS) at the National Institutes of Health. A full list of PDBP investigators can be found at https://pdbp.ninds.nih.gov/policy. The PDBP investigators have not participated in reviewing the data analysis or content of the manuscript. The Study of Isradipine as a Disease-modifying Agent in Subjects With Early Parkinson Disease, Phase 3 (STEADY-PD3) is funded by the National Institute of Neurological Disorders and Stroke (NINDS) at the National Institutes of Health with support from The Michael J. Fox Foundation and the Parkinson Study Group. For additional study information, visit https://clinicaltrials.gov/ct2/show/study/NCT02168842. The STEADY-PD3 investigators have not participated in reviewing the data analysis or content of the manuscript. Genome sequence data for the Lewy body dementia case-control cohort were generated at the Intramural Research Program of the U.S. National Institutes of Health. The study was supported in part by the National Institute on Aging (program #: 1ZIAAG000935) and the National Institute of Neurological Disorders and Stroke (program #: 1ZIANS003154).

## List of Supplemental Tables

Supplementary Table 1: QTL datasets used for analyses

Supplementary Table 2. Study population

Supplementary Table 3: Coverage details for CTSB

Supplemental Table 4: Primer and gRNA sequences used in the generation of CTSB-KO iPSC lines

Supplementary Table 5: Cell Culture Reagents and Media Compositions

Supplemental Table 6: CRISPRa and CRISPRi gRNA sequences

Supplemental Table 7: Antibodies

Supplemental Table 8: qPCR Primer sequences

Supplementary Table 9: GCTA-COLOC results

Supplementary Table 10: FINEMAP results of the nominated variants in CTSB

Supplementary Table 11: CTSB locus SMR and HEIDI analyses

Supplemental Table 12: Burden analysis for CTSB rare variants

Supplemental Table 13: Proteomics quantifications

## FIG

**Supplemental Figure 1:**
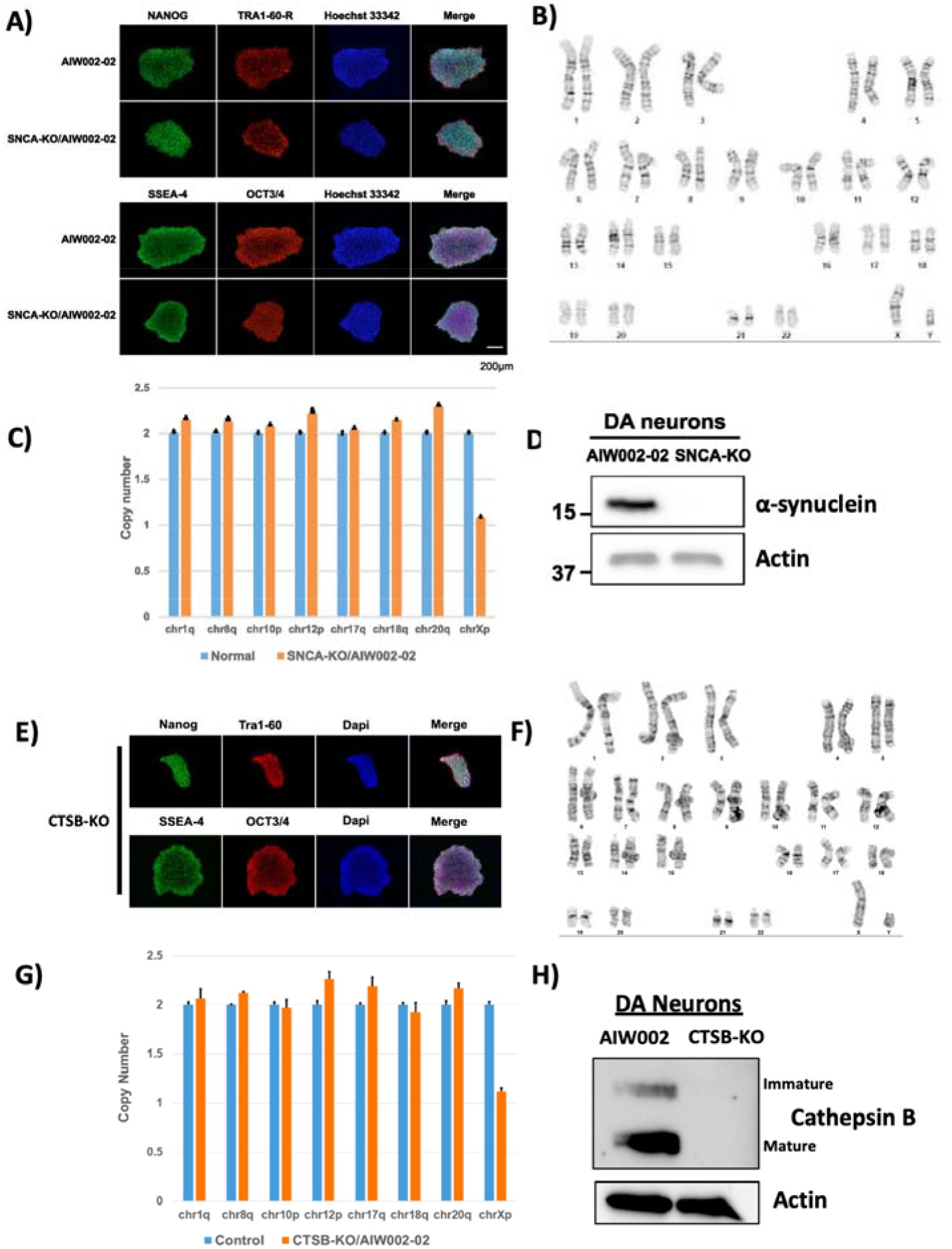
A) Immunofluorescent staining of parental (AIW002-02) and SNCA-KO iPSCs for pluripotency markers. B) Karyotype analysis of SNCA-KO iPSCs. C) Genome stability analysis of SNCA-KO iPSCs. D) Protein levels of alpha-synuclein in control versus SNCA-KO iPSCs after differentiation into dopaminergic neurons. E) Pluripotency marker expression in CTSB-KO iPSCs. F) Karyotype analysis of CTSB-KO iPSCs. G) Genome stability analysis of CTSB-KO iPSCs. H) Protein levels of cathepsin B in control versus CTSB-KO iPSCs after differentiation into dopaminergic neurons.

**Supplemental Figure 2:**
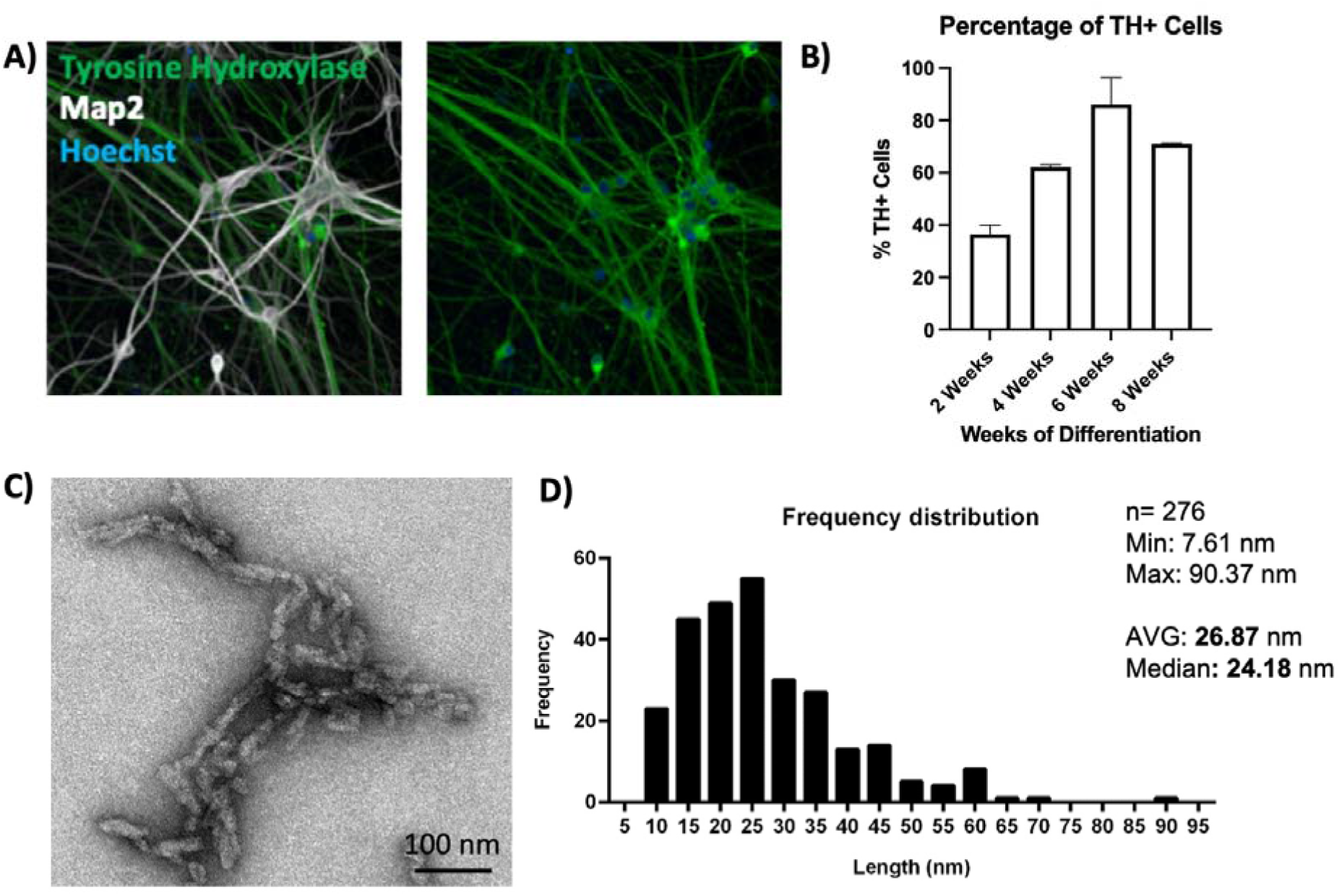
A) Differentiation of AIW002-2 iPSCs into dopaminergic neurons labelled for Map2 and tyrosine hydroxylase (TH). B) Percentage of TH-positive neurons at 2, 4, 6 and 8 weeks of differentiation. C) Representative electron microscopy imaging of α-syn PFFs. D) Measurement of α-syn PFF fibril size.

**Supplementary Figure 3:**
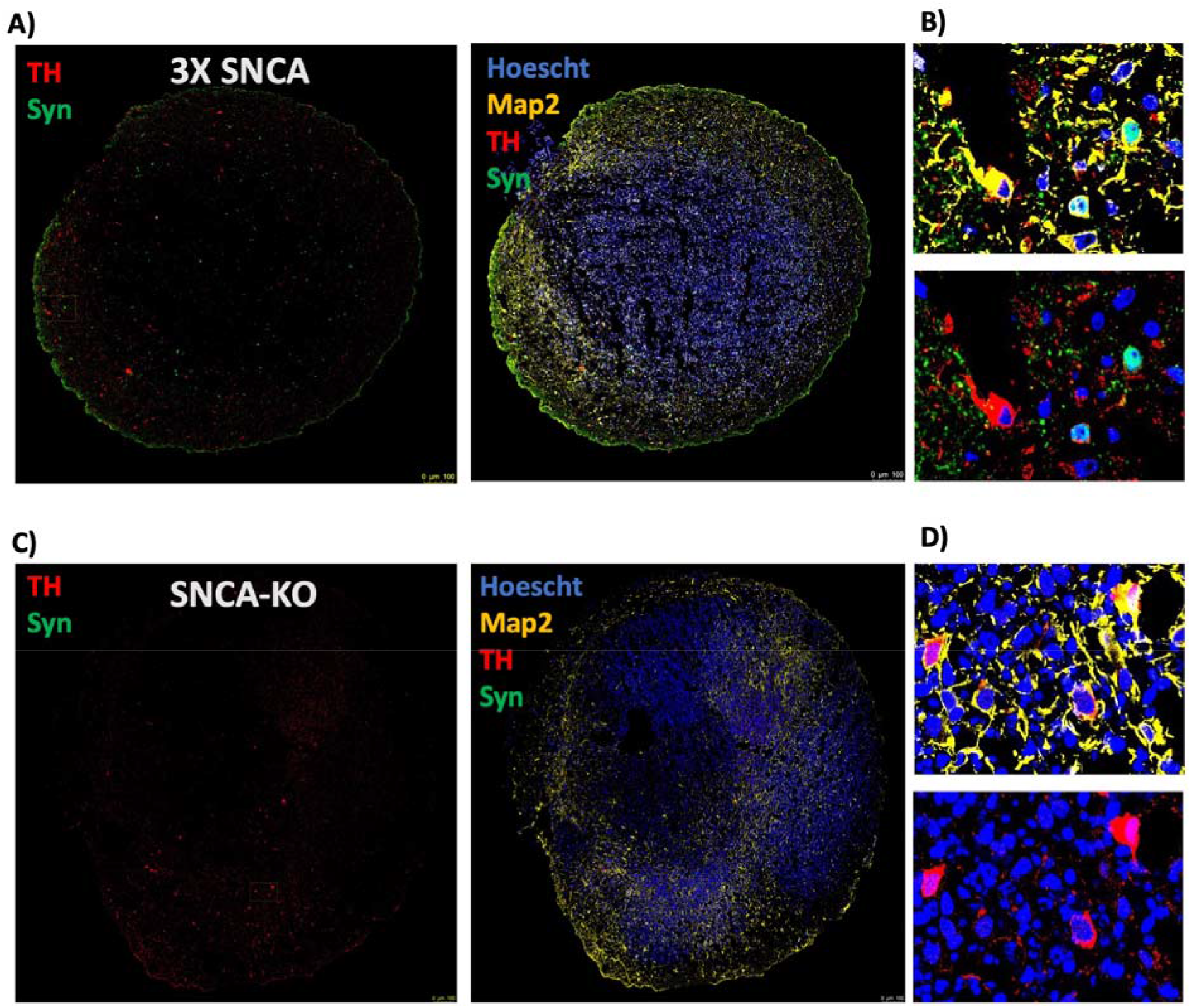
Immunofluorescent characterization of 5-month old midbrain organoids generated from SNCA triplication or SNCA-KO iPSCs. A, C) Representative immunofluorescent images depicting the entire organoid structure. B, D) Magnified images depicting individual cells. Nuclei are indicated in blue (Hoechst), total neuronal content indicated in yellow (Map2), dopaminergic neurons are stained in red (TH – tyrosine hydroxylase), and α-syn is shown in green.

**Supplemental Figure 4:**
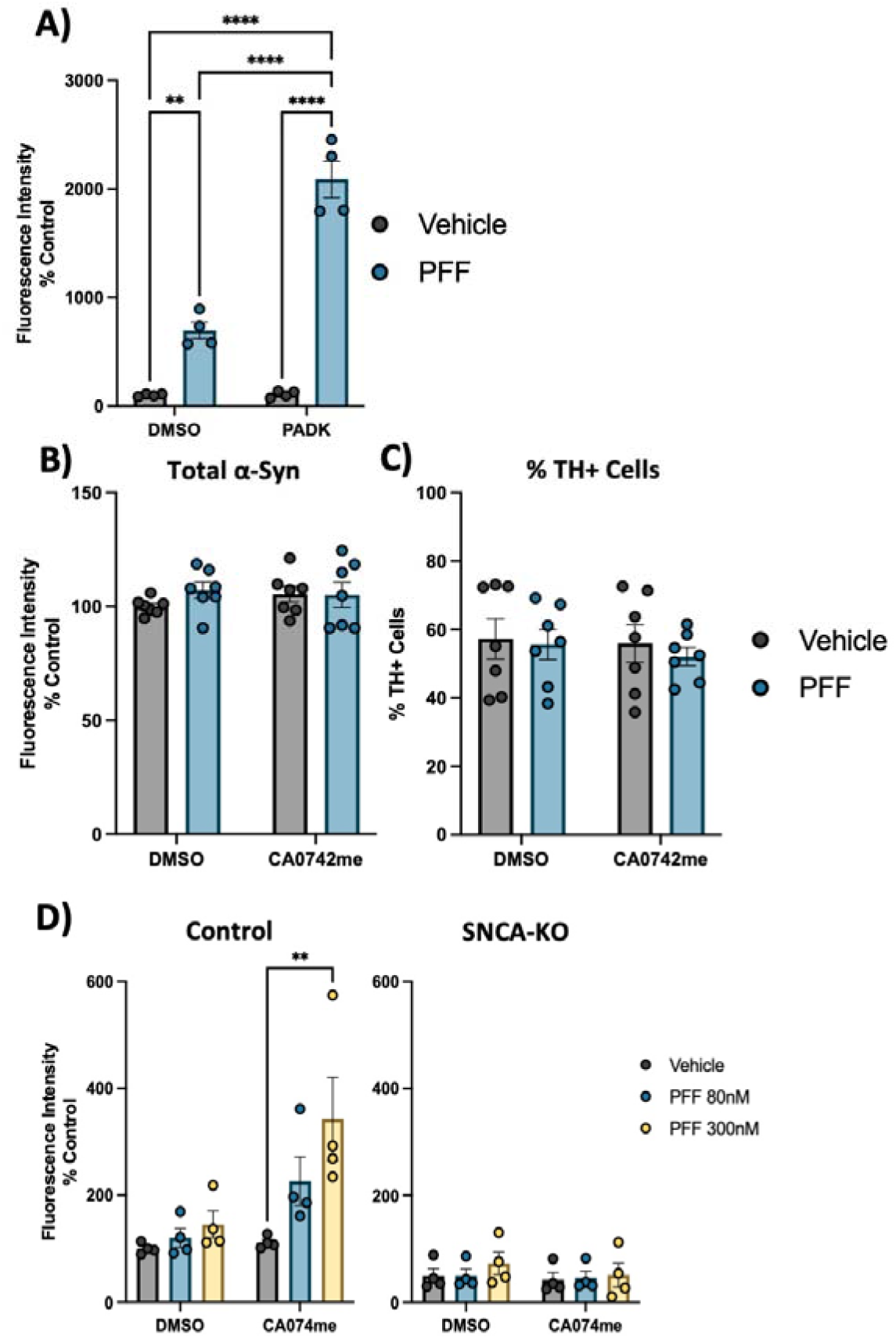
A) Quantification of pSyn-S129 in Map2-positive cells 4-weeks after exposure to PFF and/or PADK. B) Immunofluorescent quantification of total α-syn levels by high-content microscopy using the SYN1 antibody. C) Percentage of TH-positive cells in DA neuron cultures treated with CA074me (1 μM) and/or α-syn PFFs (300 nM). D) Quantification of pSyn-S129 in Map2-positive cells 3-weeks after PFF and/or CA074me treatment in either Control (AIW002-2) DA neurons or isogenic neurons lacking endogenous α-syn (AIW002-2 SNCA-KO). Bonferroni-corrected t-tests, ** p < 0.01, **** p < 0.0001.

**Supplemental Figure 5:**
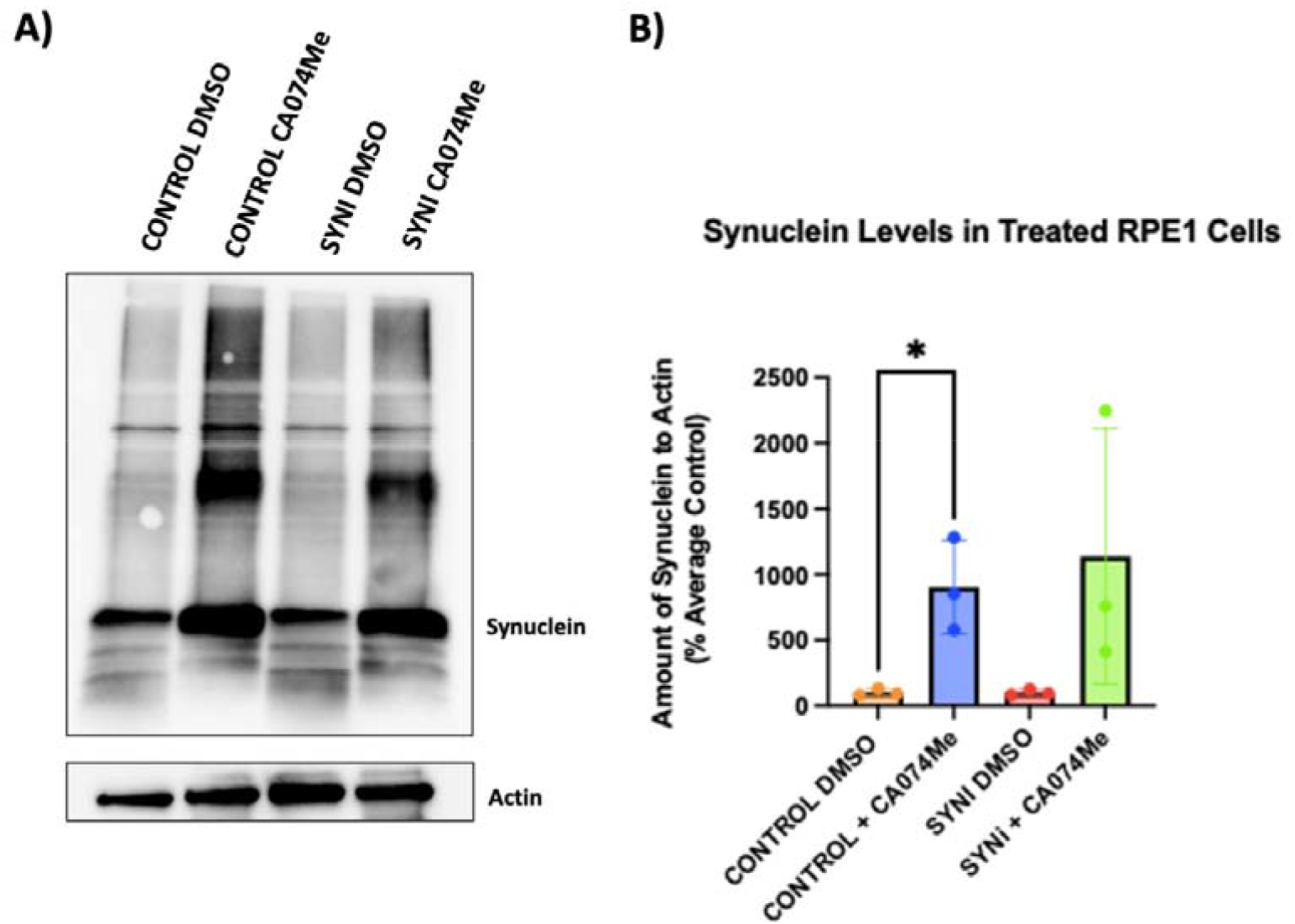
A) Representative western blot depicting protein levels of α-syn (SYN1 antibody) and actin 48-hours after treatment of RPE1 cell lines with 300nM of α-syn PFFs and/or CA074me. B) Western blot quantifications depicting levels of α-syn (SYN1 antibody – quantification of whole lane) relative to actin in PFF and/or CA074me treated RPE1 cells. T-test or Bonferroni-corrected t-tests, * p < 0.05.

**Supplemental Figure 6:**
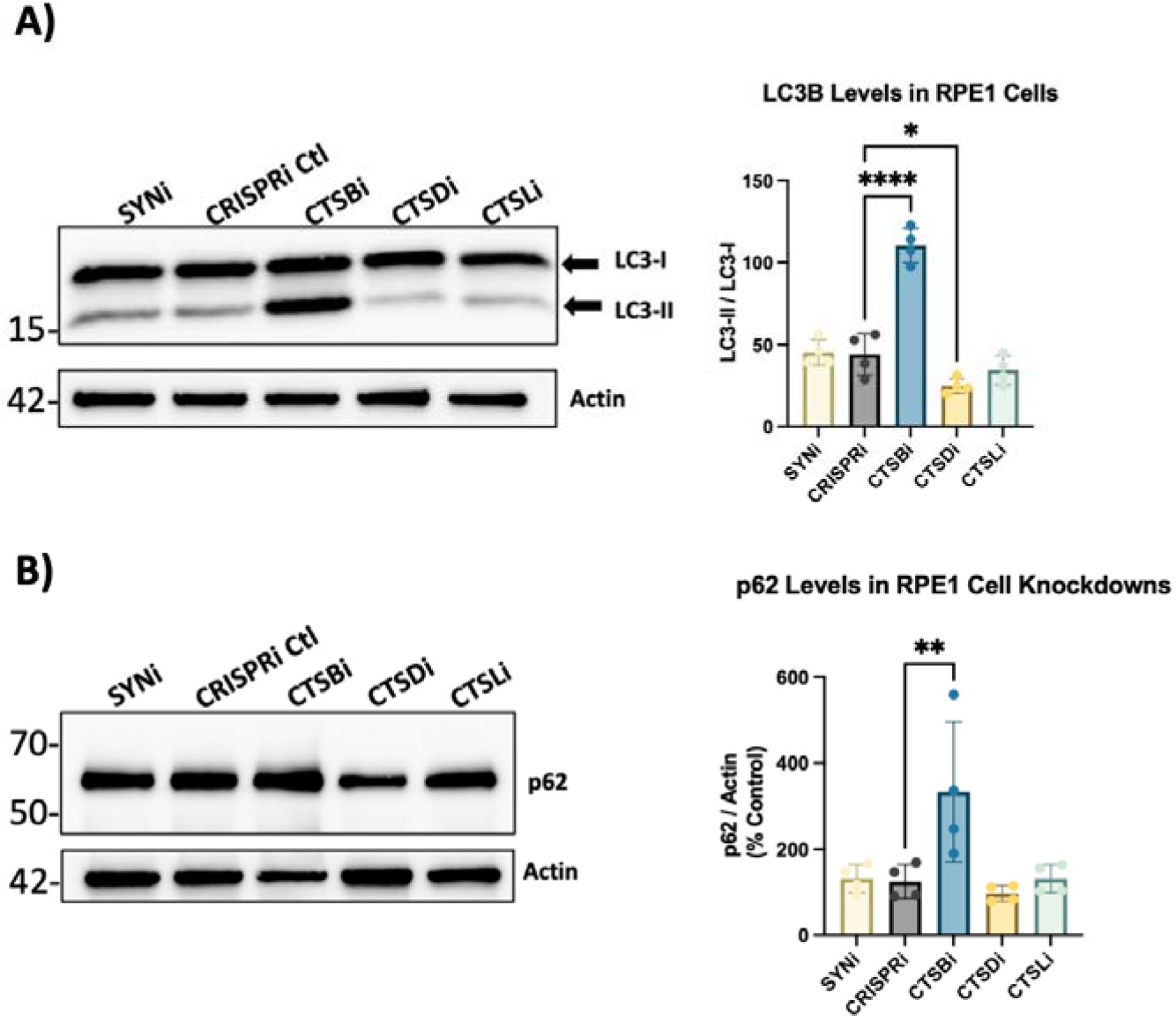
A) Representative western blots and quantification of western blots of LC3B and actin in RPE1 cell lines. B) Representative western blots and quantification of western blots of p62 and actin in RPE1 cell lines. Bonferroni-corrected t-tests, * p < 0.05, ** p < 0.01, *** p < 0.001, **** p < 0.0001.

**Supplementary Figure 7:**
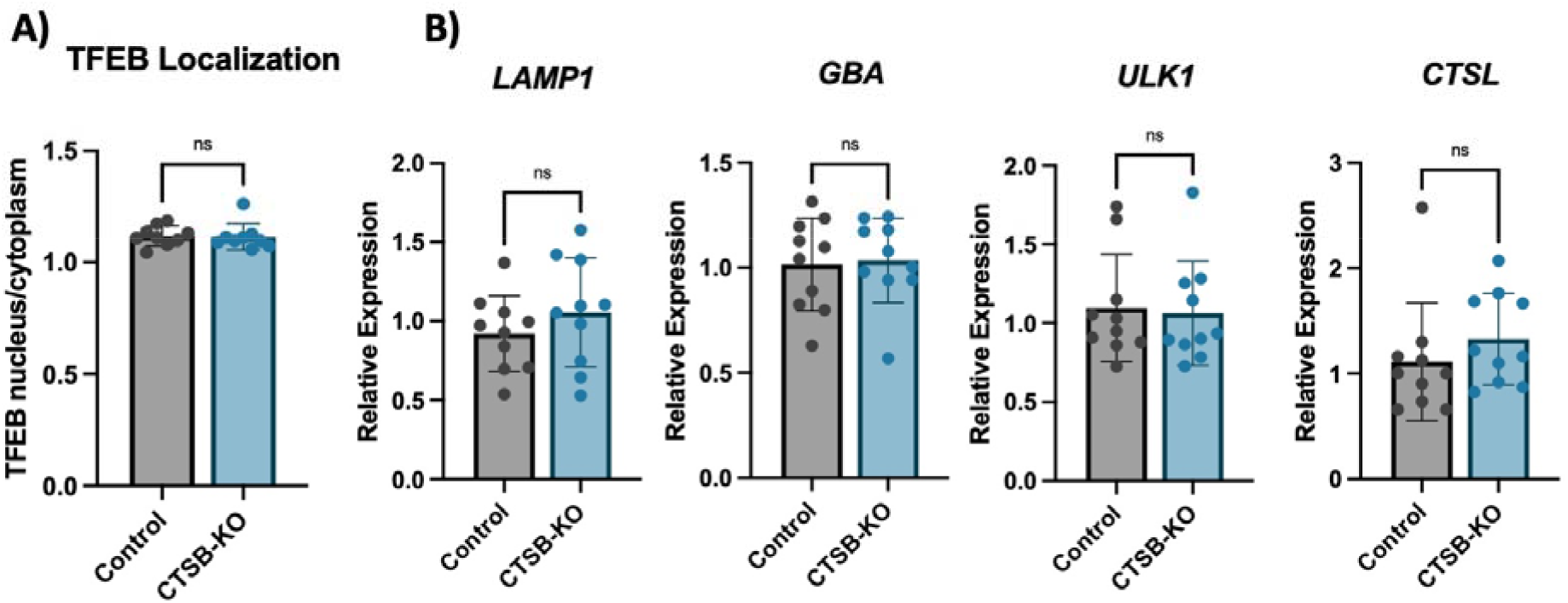
A) High-content confocal imaged-based quantification of the ratio of nuclear to cytoplasmic TFEB in iPSC-derived DA neurons based on immunofluorescence using Map2 to define the cell body area and Hoechst to define the nuclear area. B) Relative expression level of the indicated genes measured by RT-qPCR in iPSC-derived DA neurons.

**Supplemental Figure 8:**
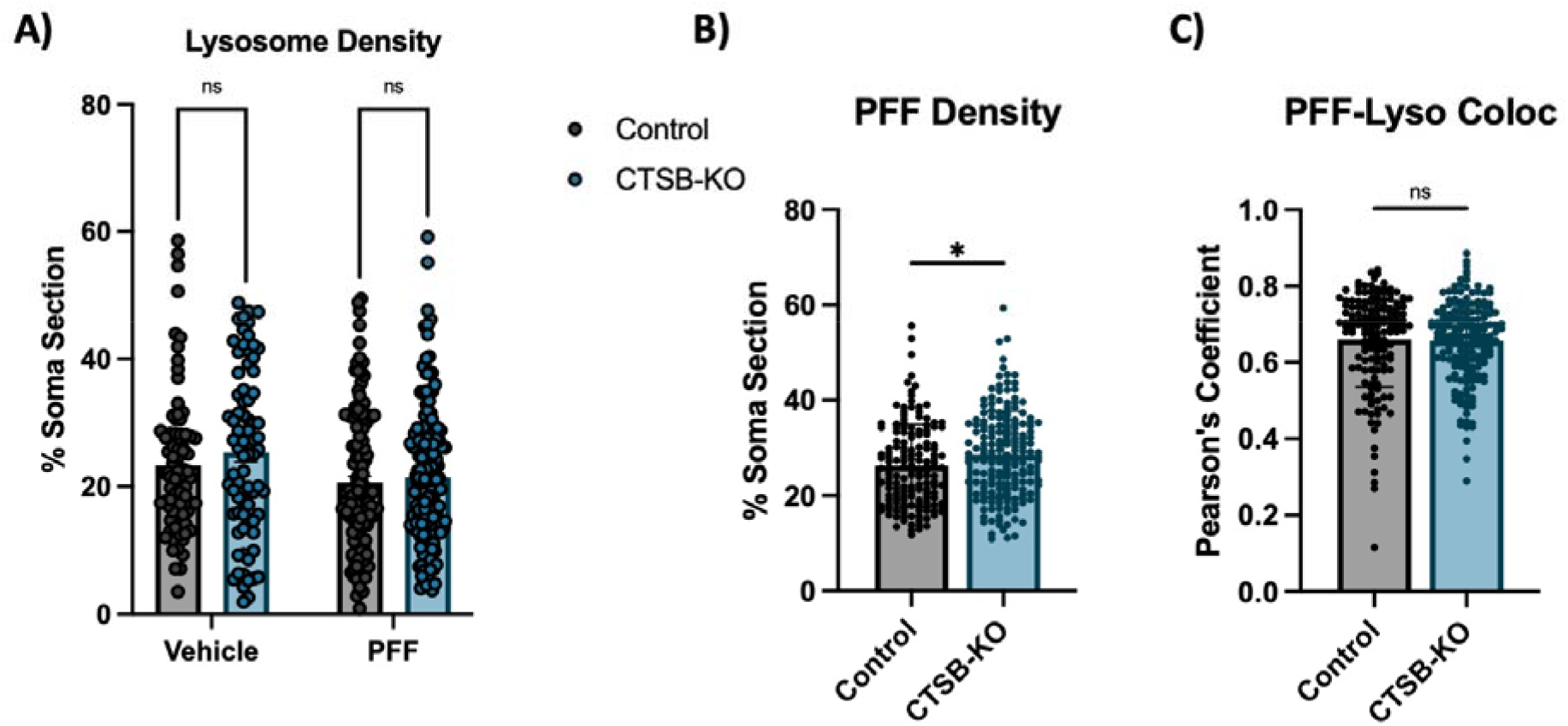
A) Quantification of lysosome density per cell body from live-cell confocal images of DA neuron cell bodies stained with lysotracker-green 72-hours after exposure to alexa-633 labelled α-syn PFFs (80 nM) and measured as the percentage of lysotracker-positive area per cell soma. B) PFF density per cell body, measured as the percentage of PFF-633-positive area per cell soma. C) Colocalization of lysotracker and PFF-633 measured using Pearson’s coefficient per cell soma. T-test, * p < 0.05, ** p < 0.01.

